# Contrasting genomic responses of hydrothermal vent animals and their symbionts to population decline after the Hunga volcanic eruption

**DOI:** 10.1101/2025.02.24.639855

**Authors:** Corinna Breusing, Michelle A. Hauer, Ian V. Hughes, Johann S. Becker, David Casagrande, Brennan T. Phillips, Peter R. Girguis, Roxanne A. Beinart

## Abstract

Genetic bottlenecks are evolutionary events that reduce the effective size and diversity of natural populations, often limiting a population’s ability to adapt to environmental change. Given the accelerating human impact on ecosystems worldwide, understanding how populations evolve after a genetic bottleneck is becoming increasingly important for species conservation. Ash deposits from the 2022 Hunga volcanic eruption in the Southwest Pacific led to a drastic decline of animal symbioses associated with hydrothermal vents in this region, allowing insights into the effects of population bottlenecks in the deep sea. Here, we applied metagenomic sequencing to pre- and post-eruption samples of mollusk-microbial symbioses from the Lau Basin to investigate patterns of genetic variation and effective population size. Our data indicate that animal host populations currently show only small changes in genome-wide diversity but in most cases experienced a long-term decline in effective size that was likely intensified by the volcanic impact. By contrast, symbiont populations exhibited a notable decrease in genomic variation, including loss of certain habitat-specific strains. However, detection of environmental symbiont sequences suggests that lost strain diversity might be recovered from the free-living symbiont pool. The differences between host and symbiont populations might be related to their contrasting genetic structures and levels of gene flow, although the full extent of population bottlenecks in the host animals might only be recognizable after a few generations. These results add to our understanding of the evolutionary dynamics of animal-microbe populations following a natural disturbance and help assess their resilience to potential future anthropogenic impacts.

## Introduction

Genetic diversity is crucial for the long-term persistence of species as it enables populations to adapt to environmental change (Kardos et al. 2021; Thompson et al. 2023). Multiple interacting factors influence the genetic variation within a population, including the type and strength of natural selection, genetic drift, the level of gene flow and the population’s effective size (N_e_), i.e., the number of individuals that are actively interbreeding (Charlesworth 2006; Reed 2007; Walsh and Blows 2009; Assis et al. 2016; Ellegren and Galtier 2016). Bottleneck effects caused by natural disasters or human activities are common evolutionary scenarios that strongly reduce the effective size and associated genetic variation of a population. Depending on their severity, genetic bottlenecks can strongly decrease fitness and adaptive potential of the affected population by limiting the efficacy of natural selection and increasing the impact of genetic drift, ultimately enhancing the risk of extinction (Whitlock 2000; Markert et al. 2010; Gamblin et al. 2024). Alternatively, selection after a bottleneck event can also favor genotypes or lineages with resilience traits, ultimately promoting rapid adaptation to the conditions of the disturbance (reviewed in Coleman and Wernberg 2020). Because of the uncertain evolutionary trajectories after a bottleneck event, information on effective population sizes and genetic diversity is increasingly considered in management plans for threatened species (Willi et al. 2021; Ottewell and Byrne 2022).

Deep-sea hydrothermal vents are enigmatic marine habitats that host unique animal and microbial communities (Van Dover 2000). Many invertebrate species in these ecosystems live in nutritional symbiosis with chemosynthetic bacteria that use energy derived from chemical reductants to convert inorganic carbon to organic matter (Sogin et al. 2020, 2021). Notably, many vent-associated taxa are classified as vulnerable or endangered on the International Union for Conservation of Nature Red List (https://www.iucnredlist.org). This is, in part, due to the island-like nature of hydrothermal vents and the restricted distribution of vent-endemic species. Accordingly, plans for commercial extraction of mineral resources from deep-sea hydrothermal vents have raised concerns about the resilience of vent-associated species and ecosystems (Levin et al. 2016; Van Dover et al. 2018; Orcutt et al. 2020; Brunner et al. 2022), highlighting the need for a better understanding of the evolutionary responses of these taxa to environmental disturbances.

In 2022, the eruption of the submarine Hunga volcano (formerly Hunga Tonga–Hunga Ha apai) caused a mass mortality event in deep-sea hydrothermal vent populations within the Lau back-arc basin of the Southwest Pacific (Beinart et al. 2024), providing an opportunity to study the impacts of genetic bottlenecks in vent-associated species and their recovery process after a natural disaster. Lau Basin vent communities are primarily composed of different co-occurring gastropods (*Alviniconcha boucheti*, *A. kojimai*, *A. strummeri, Ifremeria nautilei*) and a species of bivalve (*Bathymodiolus septemdierum*). Together they represent the foundation fauna within hydrothermal ecosystems in this region (Podowski et al. 2009, 2010). All these taxa host sulfur-oxidizing campylobacterial or gammaproteobacterial symbionts that are acquired from the environment during the animal’s larval stage and are subsequently harbored intra- or extracellularly within the animal’s gill tissue (Suzuki et al. 2006a, b; Duperron 2010; Ikuta et al. 2021). Host-symbiont associations within the Lau Basin are species-specific, with each host taxon harboring only one or two symbiont species, although symbiont strain composition can vary with local environmental conditions (Breusing et al. 2022, 2023a).

*Alviniconcha*, *Ifremeria*, and *Bathymodiolus* symbioses were abundant at hydrothermal vents along the Eastern Lau Spreading Center (ELSC) before the eruption, though *A. boucheti*, *I. nautilei* and *B. septemdierum* had already experienced a degree of population loss when the northernmost vent field Kilo Moana became inactive sometime between 2009 and 2015 (Reysenbach and Seewald 2015). The volcanic impact further decimated populations of all three mollusk genera at the vent fields nearest to the Hunga eruption, with *Alviniconcha* holobionts showing stronger population declines than *Ifremeria* or *Bathymodiolus* holobionts (Beinart et al. 2024). All three genera were completely or almost entirely extirpated from the Tow Cam vent field, which experienced significant ash deposition of up to 150 cm (Beinart et al. 2024). *Alviniconcha* was also entirely extirpated from the Tahi Moana vent field, which likewise had substantial ash deposition (12–23 cm). At the ABE vent field, where ash deposits were 8–15 cm thick, only small patches of *Alviniconcha* survived. By contrast, sparse populations of *Ifremeria* and *Bathymodiolus* remained present at both Tahi Moana and ABE. All three genera sustained seemingly normal populations at the Tu’i Malila vent field, which did not have any ash deposition (Beinart et al. 2024).

In this study, we compared pre- and post-eruption populations of *Alviniconcha*, *Ifremeria* and *Bathymodiolus* symbioses to assess initial changes in genetic diversity and effective population size in both host and symbiont populations. As one determinant of recolonization potential, we further evaluated whether free-living forms of the bacterial symbionts can still be detected in the environment.

## Material and Methods

### Sample collection, DNA extraction and metagenomic sequencing

Snail and mussel samples were collected with remotely operated vehicles from five vent fields (Kilo Moana, Tow Cam, Tahi Moana, ABE, Tu’i Malila) in the Eastern Lau Spreading Center (ELSC) during expeditions on board the R/V *Thomas G. Thompson* and R/V *Falkor* in 2009, 2016 and 2022 (Table 1). Additional samples were obtained from unimpacted hydrothermal vent communities along the Tonga Volcanic Arc during the 2016 and 2022 expeditions. For this study, these samples were only used for transcriptome reconstructions of the hosts to improve assembly quality and completeness (see below).

**Table 1.**
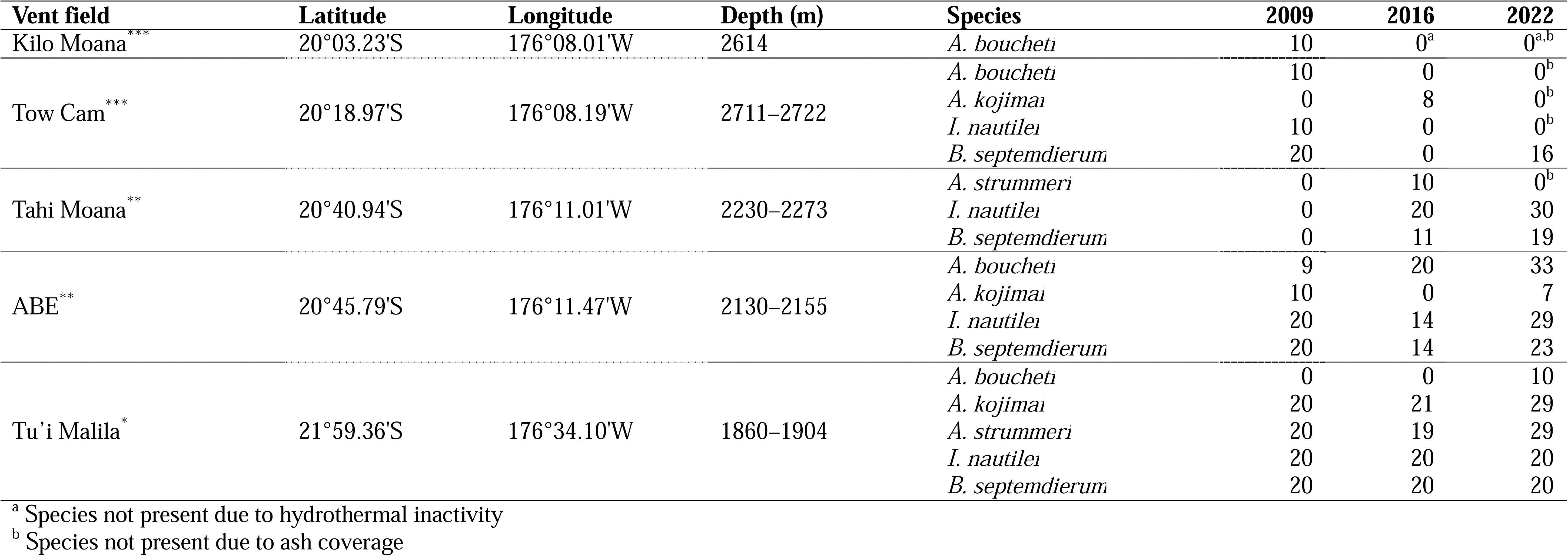
Sampling information for snail and mussel species from the Eastern Lau Spreading Center in years 2009, 2016, and 2022. Qualitative ash coverage at each vent field is indicated by stars: *** = strong coverage, ** = moderate coverage, * = little or no coverage.

All snails were collected via the remotely operated vehicles *Jason II* or *Ropos.* On board the ship, the symbiont-containing gill tissue of each animal was excised and either flash-frozen or stored in RNALater™ (Thermo Fisher Scientific, Inc., Waltham, MA, USA) at –80°C. DNA was extracted with the E.Z.N.A Mollusc DNA Kit (Omega Bio-tek, Norcross, GA) at the University of Rhode Island and sent to Psomagen, Inc. (Rockville, MD, USA) for metagenomic library preparation and sequencing. Libraries were prepared with the plexWell384 and Twist 96-Plex kits and 150 bp paired-end sequenced on NovaSeq 6000 and NovaSeq X Plus instruments to an average depth of 34,112,119 total reads per sample (Supplementary Table S1). Additional sequence data were downloaded from the National Center for Biotechnology Information (NCBI) (BioProjects: PRJNA523619, PRJNA855930) (Supplementary Table S1). Adaptors and low-quality bases were removed from the raw reads with TRIMMOMATIC (Bolger et al. 2014) before further analysis. The identity of each host species was determined morphologically and, if necessary, further verified through sequencing of the COX1 barcoding gene (Folmer et al. 1994).

### Generation of host and symbiont genetic references

To assess changes in genetic diversity in host and symbiont populations, we first assembled host transcriptomes and symbiont pangenomes as references for read mapping.

#### Host

For each *Alviniconcha* and *Ifremeria* species, total RNA of one sample from one representative vent field within the northern, southern, and central ELSC as well as the Tonga Volcanic Arc was extracted with the Zymo Direct-zol RNA Miniprep Kit (Zymo Research, Inc., Irvine, CA, USA). RNA extracts were sent to Psomagen, Inc. (Rockville, MD, USA) for library preparation with the TruSeq Stranded mRNA Kit (Illumina, Inc., San Diego, CA, USA) and 2x150 bp paired-end sequenced on a NovaSeq 6000 instrument to a depth of about 100 M reads per library. RNA-seq reads were quality-filtered, error-corrected and assembled as previously described in Breusing et al. (2022, 2023a). For *Bathymodiolus septemdierum* an existing high-quality transcriptome from Breusing et al. (2023a) was used for analysis (NCBI accession: GJZH00000000).

#### Symbiont

For each sample, filtered metagenomic reads were assembled with METASPADES (Nurk et al. 2017) using kmers from 21 to 121 and then binned into metagenome-assembled genomes (MAGs) with METABAT2 (Kang et al. 2019), MAXBIN2 (Wu et al. 216) and METAWRAP (Uritskiy et al. 2018). High-quality, non-redundant MAGs were identified with DAS TOOL (Sieber et al. 2018) and taxonomically assigned with GTDB-TK (Chaumeil et al. 2019). Additional MAGs for analysis were downloaded from NCBI BioProjects PRJNA523619 and PRJNA855930. Average nucleotide identities (ANIs) between MAGs of each bacterial taxon were computed with FASTANI (Jain et al. 2018) to determine whether they represent the same species (i.e., ANI > 95%). MAGs of each bacterial species that had CHECKM (Parks et al. 2015) completeness scores > 90% and contamination scores < 10% were used further for pangenome reconstruction with PANAROO (Tonkin-Hill et al. 2020) as in Breusing et al. (2022, 2023a). Due to the sparsity of high-quality MAGs from Kilo Moana and Tow Cam for the *A. boucheti* symbiont, completeness thresholds for MAGs from these vent fields were reduced to 80%.

### Assessment of population structure, nucleotide diversities, heterozygosity and Tajima’s D

For each host-symbiont pair we performed both field-specific and basin-wide analyses to investigate changes in genetic diversity related to the Hunga eruption. In case of the field-specific analyses, we only focused on the vent fields for each host-symbiont pair that were sampled before and after the eruption with sufficient sample coverage (i.e., >10 samples per field) and compared patterns between individual years. In case of the basin-wide analyses, we pooled all vent fields sampled for a host-symbiont pair into “Before” and “After” eruption groups. For both groups we used the same number of total samples and aimed to include at least 10 samples from each available vent field.

#### Host

Metagenomic reads of each selected sample were mapped against the corresponding host transcriptome with BOWTIE2 (Langmead & Salzberg 2012) in very sensitive local mode. The resulting BAM files were further processed with PICARD (https://github.com/broadinstitute/picard) and LOFREQ (Wilm et al. 2012) to mark optical duplicates, realign indels and recalibrate base qualities. ANGSD (Korneliussen et al. 2014) was subsequently used to estimate genotype likelihoods in each analysis group correcting for deviations from Hardy-Weinberg equilibrium and using folded site frequency spectra as prior information. Maximum and minimum depths per site were set based on the global depth distribution, while the minimum number of individuals with data to keep a site was set to 50% for the vent-specific analyses and 75% for the basin-wide analyses. For the estimation of population structure, variant sites were rigorously filtered to only include high-quality single nucleotide polymorphisms (SNPs) (-baq 1 -C 50 -minInd (nInd*0.75) -minMapQ 30 -minQ 20 - uniqueOnly 1 -remove_bads 1 -only_proper_pairs 1 -SNP_pval 1e-6 -sb_pval 0.05 -hetbias_pval 0.05 -minMaf 0.01 -skipTriallelic 1). Per-site nucleotide diversities and inter-population pairwise F_ST_s were calculated for these SNPs with ANGSD’s thetaStat and realSFS programs, respectively, while observed and expected heterozygosities were calculated from ANGSD’s HWE summary file. Tajima’s D values were inferred in 500 bp sliding windows using relaxed filter criteria to prevent bias towards positive values in the statistic (-baq 1 -C 50 -minMapQ 30 -minQ 20 - uniqueOnly 1 -remove_bads 1 -only_proper_pairs 1).

#### Symbiont

BOWTIE2 was used in very sensitive local mode to map metagenomic reads of each selected sample against the corresponding symbiont pangenome. As in the host, BAM files were further processed for removal of PCR duplicates, indel realignment and base quality recalibration. To minimize the impact of uneven read depth on variant detection, read alignments were downsampled to the lowest number of aligned reads in a sample so that per-sample pangenome coverage was at least 30X. Variants for population structure analyses were called with FREEBAYES (Garrison & Marth 2012) following Breusing et al. (2022, 2023a), excluding samples and variant sites with more than 25% of missing data. However, because most of the *A. boucheti* symbiont samples from Tu’i Malila and most of the *A. kojimai* symbiont samples from Tow Cam reached this exclusion threshold, the allowable amount of missing data was increased to 35% for the basin-wide analyses in these holobionts. Pairwise F_ST_s were calculated with the PYTHON SCIKIT-ALLEL module (https://github.com/cggh/scikit-allel), while nucleotide diversities were inferred for each vent population based on bi-allelic SNPs with the method by Romero Picazo et al. (2019). For the calculation of Tajima’s D, FREEBAYES was run without a threshold for the minimum allele frequency and the resulting VCF files were minimally filtered to remove SNP sites within a range of 5 bp around indels. Tajima’s D was estimated in windows of 500 bp with a reimplementation of VCFTOOLS (Danecek et al. 2011) that allows computation of summary statistics for haploid data (https://github.com/jydu/vcftools).

Ordination plots of variant data as well as boxplots and violin plots of nucleotide diversities, Tajima’s D and observed heterozygosities were generated in R (R Core Team 2024). Pairwise T tests or Wilcoxon rank tests were used to assess statistical differences in nucleotide diversity, Tajima’s D and observed heterozygosity among groups as well as expected and observed heterozygosity within groups. Corrections for multiple testing were performed based on the false discovery rate procedure. In the host, parameter estimates were uniformly subsampled to 100 data points per group to reduce bias related to oversampling.

### Assessment of changes in host effective population size

We used information from folded site frequency spectra based on the 2022 samples to determine if host populations had undergone a bottleneck effect following the Hunga eruption. Folded site frequency spectra for individual vent fields were computed with ANGSD’s realSFS program using the same filter criteria as for Tajima’s D. STAIRWAY PLOT2 (Liu & Fu 2020) was then used to infer changes in host effective population sizes assuming a mutation rate of 1.645e-9 per nucleotide site and generation for Mollusca and a generation time of 1 year (Allio et al. 2017).

### Detection of free-living symbionts in hydrothermal environments

To determine the potential presence of *Alviniconcha*, *Ifremeria* and *Bathymodiolus* symbionts in the environment, we sequenced the 16S V4 hypervariable region of deep-water and biofilm samples collected with a Suspended Particulate Rosette Sampler (SuPR) (Mclane Research Laboratories, Inc. Falmouth, MA USA; Breier et al. 2007) or Universal Fluid Obtainer (UFO) (National Deep Submergence Facility, Falmouth, MA, USA) and microbial colonization devices, respectively. At each vent field 15.71–47.51 L of water were pumped through 2–4 0.22 µM Express Plus Membrane filters (MilliporeSigma, MA, USA) both within close proximity to animal communities (∼10-20 cm above animal assemblages) and away from hydrothermal activity (∼10s to 100s of meters distance). A 10 µM custom nylon mesh pre-filter (Sefar Inc., NY, USA) was assembled in front of each sample filter to prevent collection of microbes associated with shed animal cells. Because the SuPR device failed during deployment at Tu’i Malila and during one dive at each ABE and Kilo Moana, 10.5 L deep water samples were additionally taken with the UFO at these localities. To account for cross-contamination using this device, empty filters were deployed but not pumped and used as negative controls. For the collection of microbial biofilms, we used weighted PVC bait cages that were enclosed by window screen mesh on either end and contained vertically held, open-ended 2-mL polypropylene tubes packed with crushed mineral substrates (andesite, basalt, glass beads or *Alviniconcha* shells) for microbial colonization. Three of these cages were deployed during the first dive at each vent field and recovered approximately two weeks later. On board ship, water and biofilm samples were preserved in RNALater™ (Thermo Fisher Scientific, Inc., Waltham, MA, USA) and frozen at –80°C. DNA from filter samples was extracted with a phenol:chloroform protocol (Supplementary Methods), while DNA from biofilms was extracted with the E.Z.N.A. Soil DNA Kit (Omega Bio-Tek, Norcross, GA, USA), including an additional 45 second bead beating step prior to proteinase K digestion. For the SuPR and UFO samples the 3 Strain Tagged Genomic DNA Even Mix (ATCC, VA, USA) was added as spike-in control and dilutions of the spike-in without DNA extract were prepared to assess cross-contamination during sample processing in the laboratory. Absolute microbial abundance data leveraging information from these spike-ins will be reported in a different study (Hauer et al. in prep). Final extracts were sent to Argonne National Laboratory (Lemont, IL, USA) for amplicon library preparation with the 515F/806R primer pairs (Apprill et al. 2015; Parada et al. 2016). All libraries were 2x150 bp paired-end sequenced to an average depth of 1,455,454 total reads/sample on a NextSeq2000 instrument. Raw reads were quality-checked with FASTQC (https://github.com/s-andrews/FastQC) and then processed with FASTP (Chen et al. 2018) and TRIMMOMATIC (Bolger et al. 2014) to remove polyG tails and adaptors, respectively. The USEARCH (Edgar 2010) and VSEARCH (Rognes et al. 2016) tools were then used to merge (-fastq_minmergelen 230 -fastq_maxmergelen 270 - fastq_maxdiffs 10 -fastq_pctid 80), orient and filter (--fastq_qmax 42 --fastq_minlen 230 -- fastq_truncqual 20 --fastq_maxee 1) the reads. Filtered sequences were denoised and decomposed into amplicon sequence variants (ASVs) and their abundances assessed via the VSEARCH --search_exact command. The R packages PHYLOSEQ (McMurdie and Holmes 2013) and METAGMISC (https://github.com/vmikk/metagMisc) were used to remove singletons as well as spurious ASVs with less than 10 average total read counts per sample and a prevalence of less than 5%. ASV sequences were imported into QIIME2 (https://qiime2.org) for taxonomic classification with a Naive Bayes classifiers trained on the 515F/806R primer region and considering taxonomic weights specific to seawater and marine biofilms (Kaehler et al. 2019). Potential symbiont ASVs from the classes Gammaproteobacteria and Campylobacteria were blasted against a reference database of verified *Alviniconcha*, *Ifremeria* and *Bathymodiolus* 16S rRNA symbiont sequences. ASVs with greater than 99% identity to a reference sequence were considered free-living symbionts. Read counts from all samples were corrected by the maximum amount of corresponding ASV reads in the spike-in control samples. Because symbiont sequences were abundant in the UFO control filters except at the Tu’i Malila vent field, the Kilo Moana and ABE UFO samples were not further considered for the calculation of overall symbiont read counts. In the case of Tu’i Malila, the UFO samples were further corrected by the read counts in the UFO negative control. Samples with less than 20,000 corrected read counts were excluded from further analyses.

## Results

### Host-symbiont population genetic structure

Co-assemblies of three to five gill RNA samples for each *Alviniconcha* and *Ifremeria* species resulted in 36,164–57,396 transcripts (63.91–82.73 Mb) that represented 75.80–81.40% of the host transcriptomes (Supplementary Table S2). Read mapping against these reference transcripts and the previously reconstructed *B. septemdierum* transcriptome (Breusing et al. 2023a) recovered 14,372–70,374 high-quality SNPs for population genetic analysis. For the three *Alviniconcha* species, principal component plots based on these SNPs showed structuring of the northern and central vent fields (i.e., Kilo Moana, Tow Cam, Tahi Moana) relative to the southern fields (i.e., Tu’i Malila) (Fig. 1). Pairwise F_ST_ values indicated an increasing isolation with distance (Supplementary Table S3), with highest differentiation being observed between the Kilo Moana and Tow Cam populations and the newly discovered Tu’i Malila population of *A. boucheti* (0.1147–0.1209). In general, however estimates for all other vent field comparisons were modest (0.0334–0.0770) in accordance with previous studies (Breusing et al. 2022). In *Ifremeria* and *Bathymodiolus*, no geographic pattern could be observed (Fig. 1) and F_ST_ indices for the same population comparisons were generally smaller than in *Alviniconcha*, with *Bathymodiolus* exhibiting the least amount of differentiation (*Ifremeria*: 0.0162–0.0505; *Bathymodiolus*: 0.0092–0.0141). Overall, no genetic differences could be observed between populations of the same vent fields before and after the eruption, except for *A. kojimai* where the pre- and post-eruption populations at ABE appeared to form separate clusters.

**Fig. 1.**
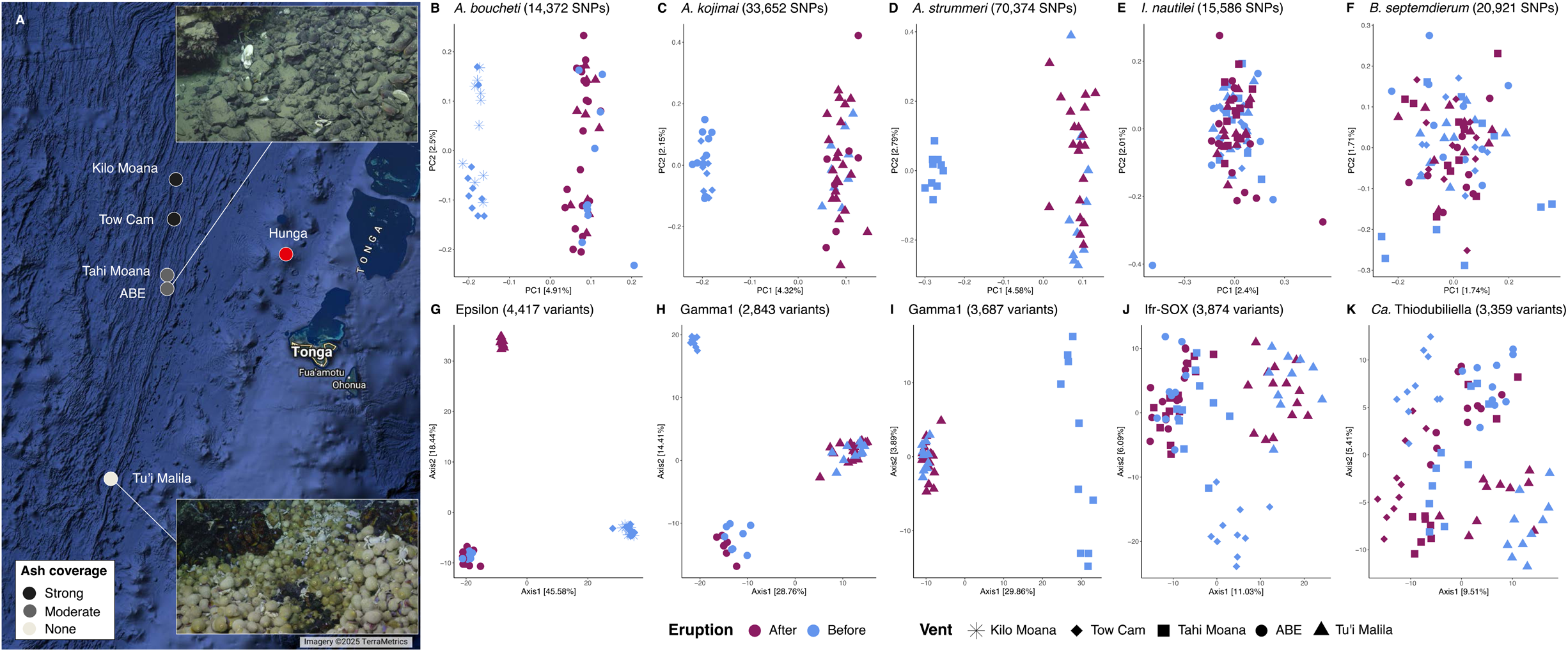
Geographic map of vent field locations (A) and principal component plots for the *Alviniconcha*, *Ifremeria* and *Bathymodiolus* host species (B–F) and their corresponding symbionts (G–K) based on genomic variant data. Photo insets in panel (A) show healthy mollusk communities at Tu’i Malila (bottom) and ash-covered mollusk communities at ABE (top). Blue colors in the PCA plots indicate samples collected pre-eruption, while red colors denote samples collected post-eruption. Map data ©2025 Google.

In the symbiont, 79–172 high-quality MAGs were used to create ∼2.87–4.61 Mb pangenomes for each bacterial species as references for population genetic analyses (Supplementary Tables S4, S5). After read mapping against these pangenomes, 2,843–4,417 high-quality variants were obtained. In agreement with previous reports (Breusing et al. 2022) and in contrast to patterns in the host, principal coordinate plots and pairwise F_ST_ calculations based on these variants indicated strong subdivision of *Alviniconcha* symbiont populations between vent localities (F_ST_: 0.1487–0.8617). Subdivision of the *Ifremeria* (F_ST_: 0.0144–0.2504) and *Bathymodiolus* (F_ST_: 0.0507–0.2122) symbiont populations was present though weaker than in the *Alviniconcha* symbionts, with the *Bathymodiolus Ca.* Thiodubiliella symbiont showing the least amount of structure (Fig. 1; Supplementary Table S3; Breusing et al. 2023a). The *Alviniconcha* and *Ifremeria* symbiont strains from the northernmost vent fields were absent after the eruption, while no change was observed for the *Bathymodiolus* symbiont strains.

### Changes in genetic diversity

With few exceptions, nucleotide diversities and mean observed heterozygosities calculated for high-quality variant sites did not differ significantly between pre- and post-eruption years in any host species in either the basin-wide or vent-specific analyses (Fig. 2–3, S1–S2). Furthermore, mean observed heterozygosities did not consistently deviate from expectations in the post-compared to pre-eruption years, except in *B. septemdierum*, where almost all comparisons indicated a significant heterozygote excess in the post-eruption populations (Fig. 3, S2). Significant heterozygote excess was also observed in a few pre-eruption populations of *B. septemdierum* (2009: Tow Cam; 2016: Tahi Moana, ABE) as well as the ABE 2016 and 2022 populations of *I. nautilei*, while significant heterozygote deficiencies were observed in the ABE 2009 population of *A. boucheti* and in the Tu’i Malila 2009 population of *A. strummeri* (Fig. S2).

**Fig. 2.**
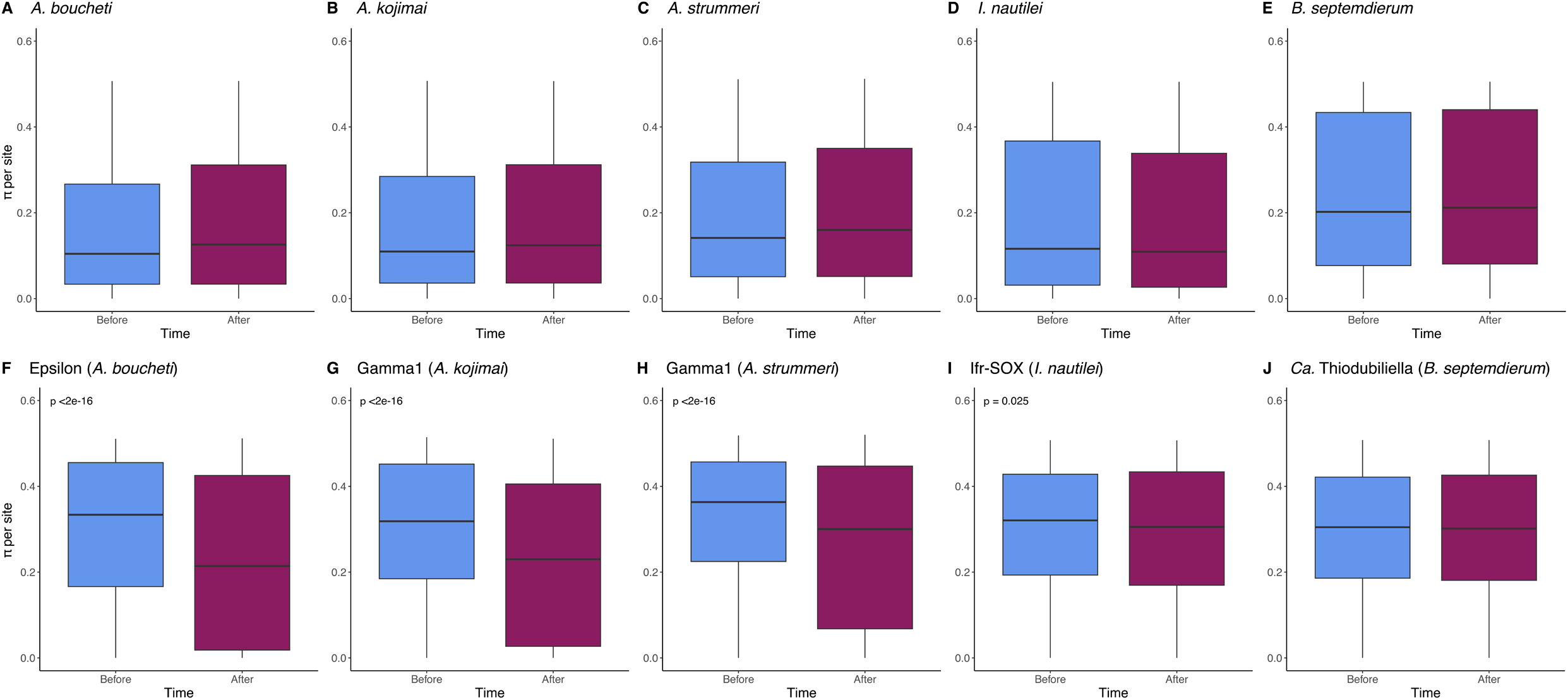
Bar plots of per-site nucleotide diversities for high-quality variants in host (A–E) and symbiont (F–J) populations before and after the eruption. The plots show results for pre- and post-eruption population pools across the Lau Basin.

**Fig. 3.**
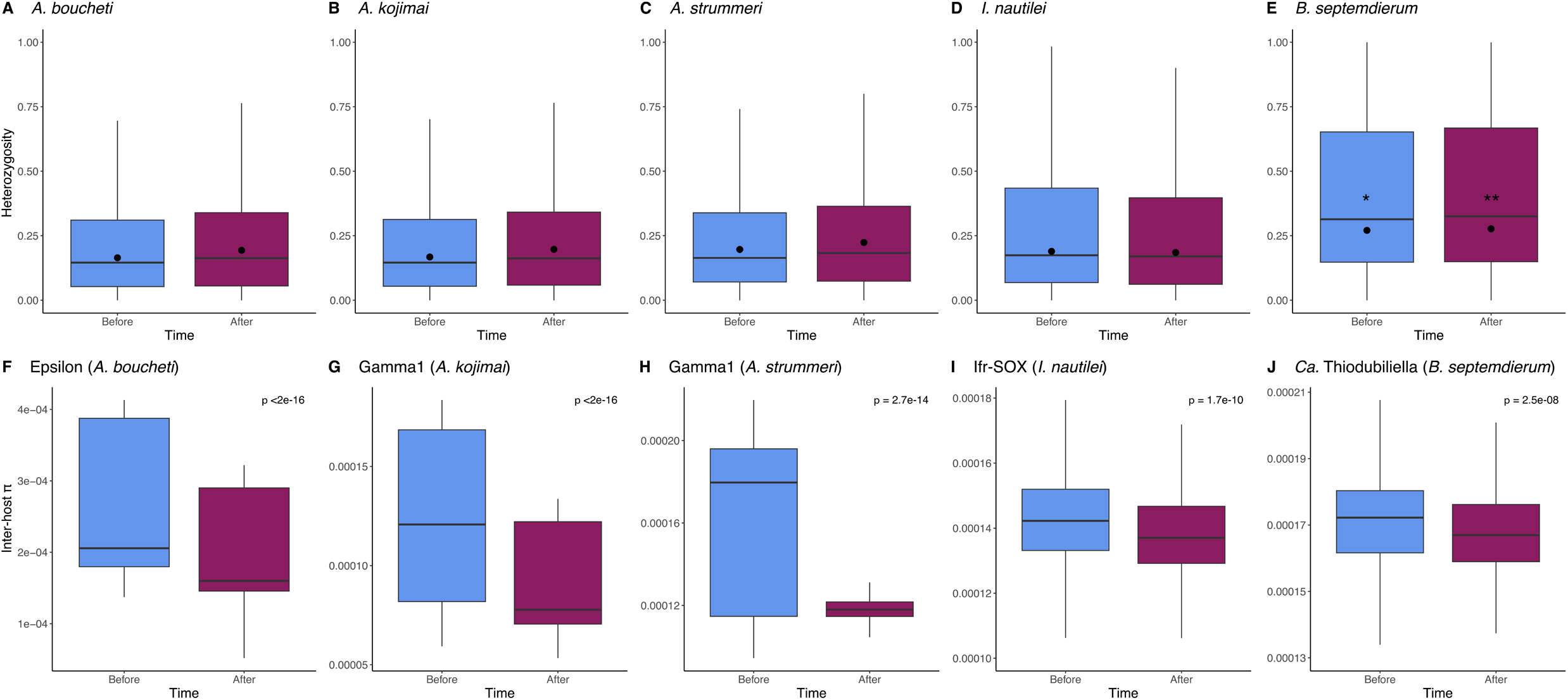
Bar plots of observed heterozygosities at high-quality variant sites in host populations (A– E) and inter-host diversities between symbiont populations (F–J) before and after the eruption. Black dots in panels A–E show the median expected heterozygosities, with stars within bar plots indicate significant differences between expected and observed mean heterozygosities for a group. * *p* < 0.05, ** *p* < 0.01, *** *p* < 0.001

In the symbiont populations, genetic diversity showed a significant decline after the eruption in the basin-wide analysis, both when looking at overall genomic variation (exception: *Ca.* Thiodubiliella) and variation between individual hosts (Fig. 2, 3). Similar to patterns in the host, however, no consistent trend of decreasing nucleotide diversity post-eruption was observed for the year-to-year comparisons of individual vent locations (Fig. S3, S4). In these comparisons per-site and inter-host nucleotide diversities were typically either unchanged or in some cases increased in 2022 compared to 2009 or 2016 (Fig. S3, S4).

### Bottleneck effects and changes in effective population size

The distribution of Tajima’s D values in the hosts suggested predominantly neutrally evolving populations without strong evidence for recent allele frequency changes (Fig. 4, S5). In general, no significant differences in Tajima’s D could be found between the pre- and post-eruption populations of any host species. Notable exceptions were the basin-wide comparisons in *I. nautilei* and *B. septemdierum* and the between-year comparisons in the ABE populations of *A. boucheti* and *I. nautilei* where Tajima’s D values were slightly more negative after the eruption, suggesting a potential onset of population expansion following a recent bottleneck (Fig. 4). By contrast, assessment of changes in effective population size through stairway plots indicated a longer-term decline for most host populations, except for the Tu’i Malila populations of *A. strummeri* and the ABE population of *A. boucheti* (Fig. 5).

**Fig. 4.**
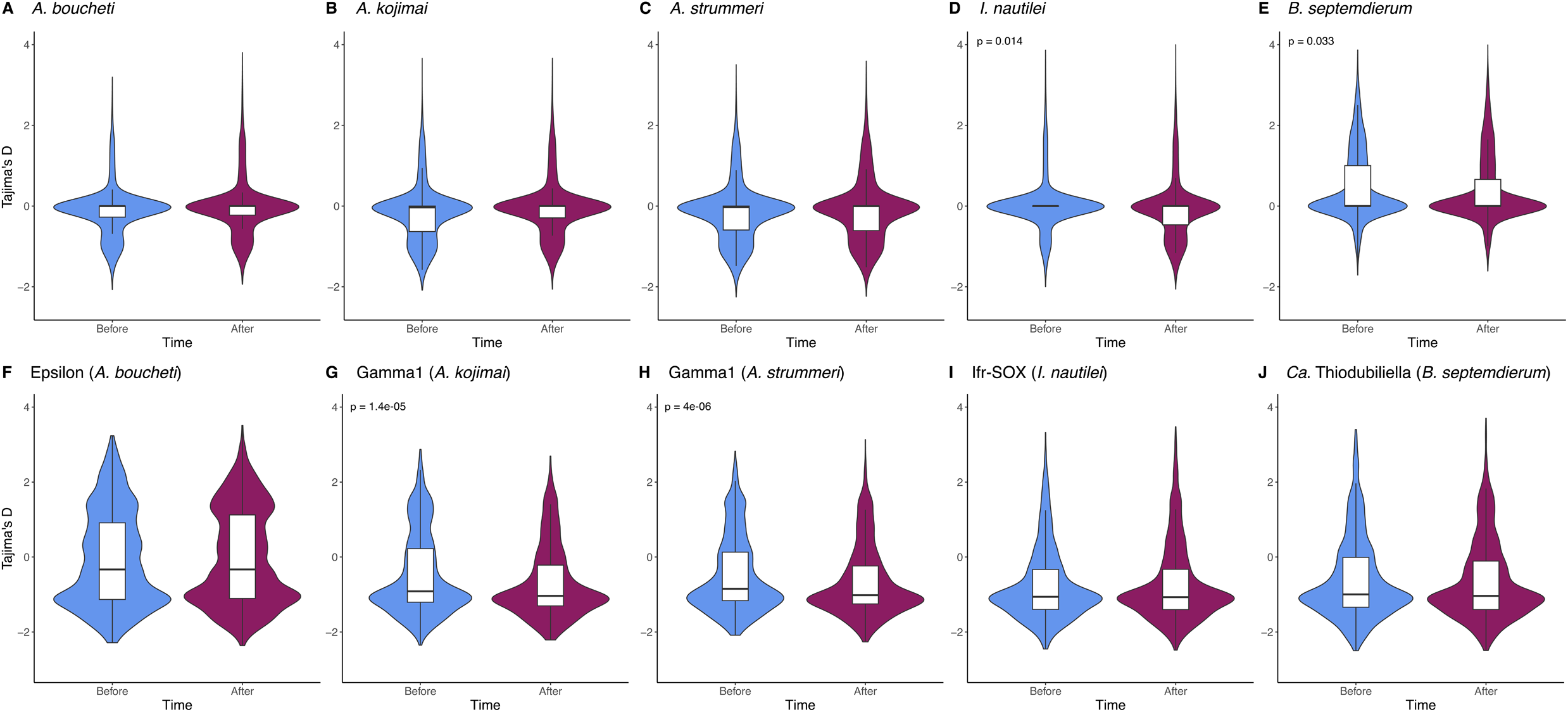
Violin plots with integrated bar plots of Tajima’s D values calculated across the host transcriptomes (A–F) and symbiont pangenomes (F–J).

**Fig. 5.**
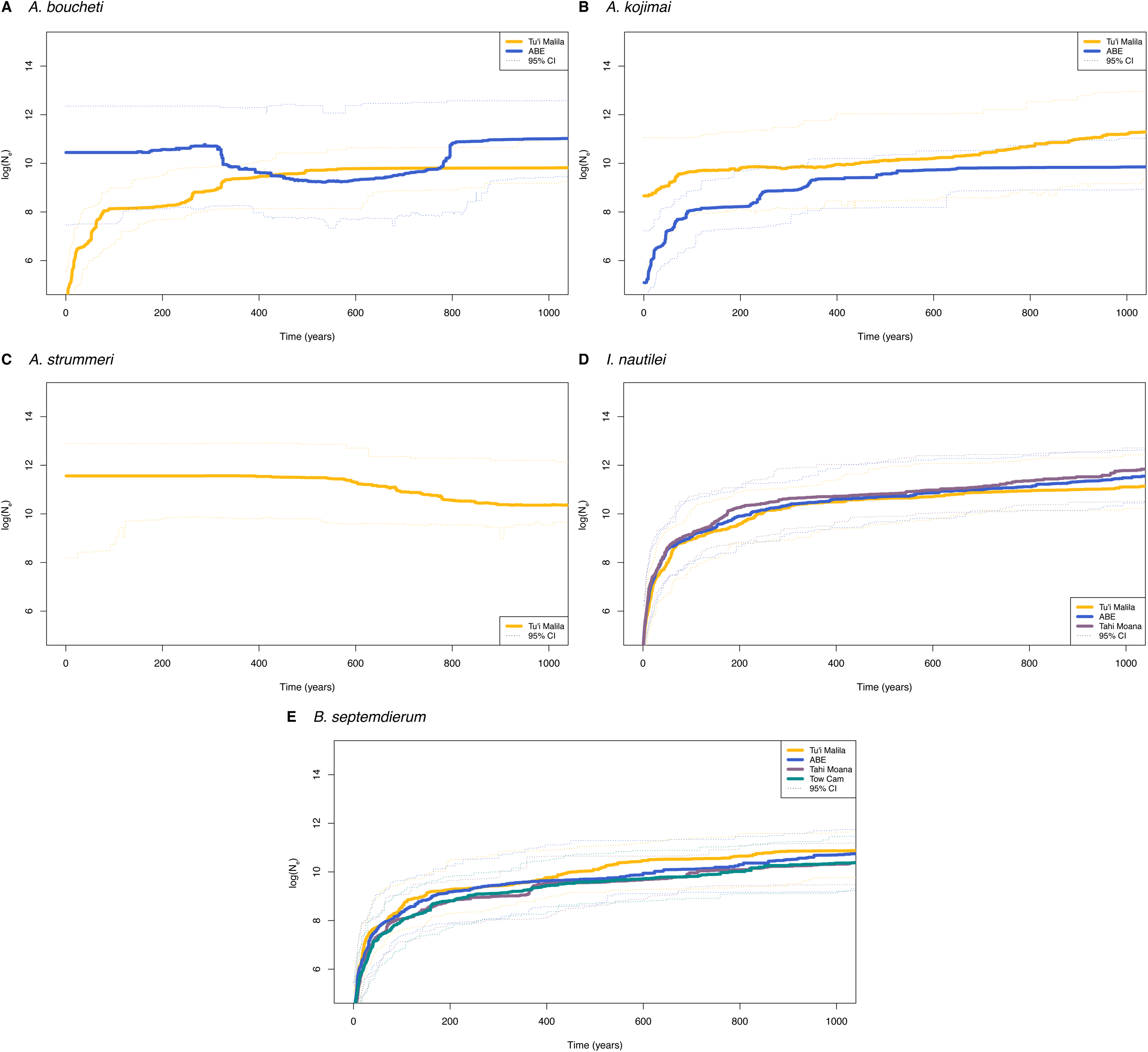
Stairway plots showing changes in host effective population sizes over the last 1000 years.

In the symbionts, Tajima’s D statistics were overall shifted to negative values, indicative of populations experiencing positive selection (Fig. 4, S6). In the basin-wide comparisons, only the Gamma1 symbiont of *A. kojimai* and *A. strummeri* showed a significant decrease in Tajima’s D following the eruption (Fig. 4). In the vent-specific comparisons, by contrast, almost all post-eruption symbiont populations indicated a slight but significant drop in Tajima’s D (Fig. S6).

### Presence of free-living symbionts in the environment

Amplicon denoising and filtering resulted in 7,660 final ASVs. Seventeen of these ASVs had significant similarity to *Alviniconcha* (*Sulfurimonas*: 5, Gamma1-like: 6), *Ifremeria* (*Thiolapillus*: 1) and *Bathymodiolus* (*Ca.* Thiodubiliella: 5) symbiont species (Supplementary Table S6). Relative abundances for these environmental symbiont ASVs varied notably with sample type (biofilm *versus* deep water) and location (Figure 6; Supplementary Table S7). The *Ca.* Thiodubiliella symbiont was abundant at all sampled vent fields, with highest average proportions being observed in seawater followed by hydrothermal fluids and no read counts being observed in biofilm samples (Figure 6; Supplementary Table S7). By contrast, 16S rRNA sequences of the snail symbionts were predominantly recovered from biofilms, with the *Sulfurimonas* symbionts being mainly observed at the central vent fields (Tahi Moana and ABE), the Gamma1-like symbionts being mainly observed at the southernmost vent field Tu’i Malila and the *Thiolapillus* symbiont showing low but similar proportions across all localities (Figure 6; Supplementary Table S7). Within hydrothermal fluids, the highest proportions for the Gamma1-like and *Thiolapillus* symbionts were detected at Tu’i Malila and Tahi Moana, respectively, while proportions for the *Sulfurimonas* symbionts increased from south to north (Figure 6; Supplementary Table S7). Relative abundances for the snail symbionts were overall lowest in seawater samples away from hydrothermal activity and at the extinct vent field Kilo Moana (Figure 6; Supplementary Table S7). Across all samples, the highest number of reads was recovered for the *Ca.* Thiodubiliella symbionts, whereas the lowest number of reads was detected for the *Thiolapillus* and Gamma1-like symbionts (Supplementary Table S7).

**Fig. 6.**
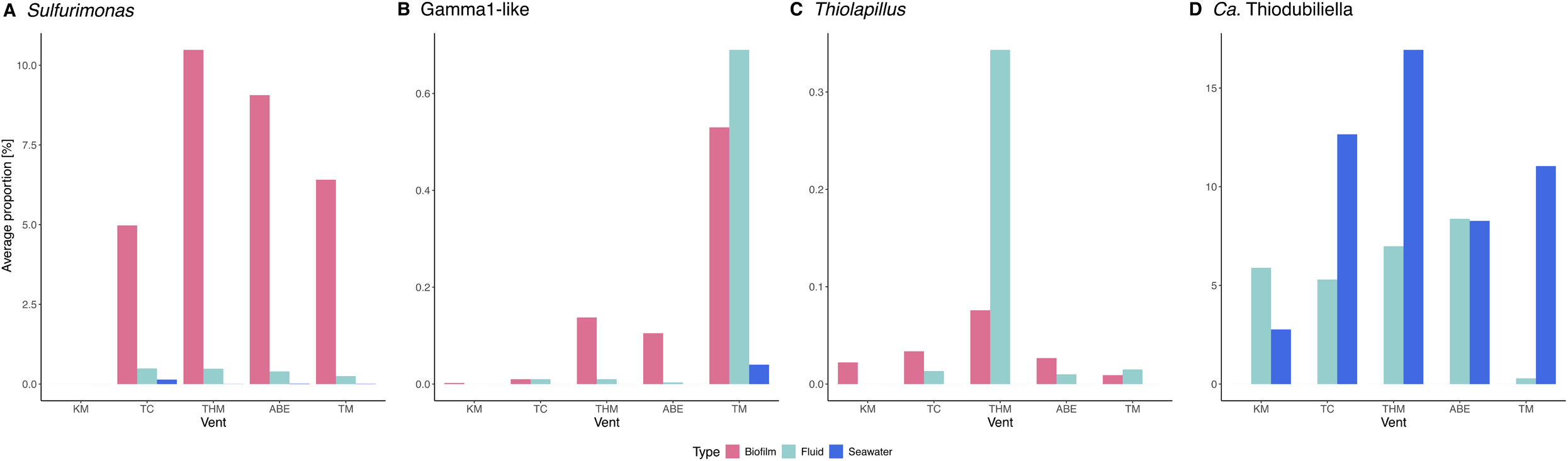
Average relative abundances of free-living symbionts on mineral surfaces (biofilms), in hydrothermal fluids near animal/shell patches and away from hydrothermal activity (seawater).

## Discussion

Natural populations often experience changes in effective population size and genetic variation due to bottleneck effects caused by severe environmental disturbances. The long-term consequences of bottleneck events are usually difficult to predict as observations from many wild populations are missing and insights from one system might not be extrapolatable to another because evolutionary trajectories depend on the demographic and life-history traits of a species (Hoelzel 1999; Olazcuaga et al. 2023; Gamblin et al. 2024). In this study we aimed to characterize changes in genetic diversity and effective population size of Western Pacific hydrothermal vent symbioses that were recently impacted by the eruption of the Hunga submarine volcano.

Theory predicts that bottleneck effects are strongest in small populations with low gene flow (Star and Spencer 2013), while immigration from other populations can neutralize the resulting loss of genetic diversity (Vilá et al. 2003; Jangjoo et al. 2016; Yu et al. 2020). The population genetic patterns found for the animal host species follow these predictions. No immediate changes in nucleotide diversities or consistent deviations from expected heterozygosities could be observed, which might be related to the high gene flow reported for these populations (Breusing et al. 2022, 2023a, b), although a couple of generations might have to pass to determine how genetic variation will be affected in the host populations. Nevertheless, at least for the *Alviniconcha* host species our data revealed the presence of population genetic structure contrary to our previous work (Breusing et al. 2022), indicating that some genotypic variation was lost in these taxa even if other measures of diversity were unchanged. In addition, almost all post-eruption populations of *B. septemdierum* indicated an excess of heterozygotes, which is a typical pattern found after recent genetic bottlenecks given that allelic diversity decreases more quickly than heterozygosity (Cornuet and Luikart 1996). In accordance with these findings, most host populations showed an ongoing decline in effective population size that intensified in the recent past. As estimates for generation times and mutation rates are unknown for the studied deep-sea organisms, absolute values from these results must be interpreted with caution. For example, slower mutation rates would result in a more modest decrease in effective population size and higher absolute estimates for N_e_. Assuming that the overall patterns are reliable, however, our findings indicate that the observed decrease in N_e_ seemed to have started before the recent volcanic eruption, suggesting that vent populations in the Lau Basin might have experienced a longer-term change in hydrothermal activity, as exemplified by the extinction of the Kilo Moana vent field between 2009 and 2015 (Reysenbach & Seewald 2015). Alternatively, it is possible that the demographic patterns in the studied deep-sea populations reflect a general trend for marine animal species, which are increasingly impacted by climate change even in the remote recesses of the deep ocean (Levin and Le Bris 2015; Penn and Deutsch 2022).

Opposite to the patterns seen in the animal hosts, symbiont populations associated with *Alviniconcha* and *Ifremeria* showed a significant reduction in genomic diversity, at least on a basin-wide scale. Our previous work demonstrated that symbiont strains of these host species are locally adapted to the conditions at their respective vent fields (Breusing et al. 2022), leading to strong population differentiation among localities. Because *Alviniconcha* and *Ifremeria* populations at the northern part of the Lau Basin were mostly eradicated, the resulting loss of vent-specific symbiont strains should inevitably reduce genetic diversity in the symbiont meta-population for these holobionts. A decline in basin-wide genetic variation was not observed for the *Bathymodiolus* symbiont, even though inter-host diversity was slightly reduced, implying that at least some minority strains were lost. These differences can likely be explained by the fact that the mussel populations were not as impacted by the eruption as the two gastropod genera and remained present even at the most affected northern vent fields. In addition, *Bathymodiolus* symbiont strains are much less structured across geographic locations compared to the *Alviniconcha* and *Ifremeria* symbionts due to reduced selective pressures (Breusing et al. 2023a), which should lower the chances of basin-wide strain loss during environmental disturbances. Within a given vent field, symbiont genetic diversity was generally unchanged among pre- and post-eruption years for all holobionts. This could partly be related to the fact that we needed to focus these analyses on less impacted vent populations for which sufficient post-eruption samples could be obtained, especially in the case of the *Alviniconcha* symbioses. On the other hand, it is possible that within a given vent field symbiont strains are distributed widely enough among different host individuals to prevent loss of local symbiont genetic variation even when host populations are diminished.

Although our data suggest that certain symbiont strains vanished from their hosts after the eruption, it is possible that these strains are still present in the vent environment in their free-living forms. We have some preliminary support for this assumption as environmental 16S rRNA sequences of *Alviniconcha, Ifremeria* and *Bathymodiolus* symbionts were in fact found at all investigated hydrothermal vent fields. However, because the 16S V4 hypervariable region does not have enough resolution to assign strain identities to the recovered sequences (Edgar 2018), we are currently unable to determine if they comprise some of the same strains that were also found in the hosts or completely different strains that are part of the free-living symbiont pool. Further metagenomic sequencing of the environmental samples taken in this study will be necessary to answer this question.

Apart from implying a route for replenishment of lost strain-level diversity, our data provide some first insights into where free-living vent mollusk symbionts are distributed in the environment. Previous studies of *Riftia pachyptila* tubeworms and *Bathymodiolus septemdierum* mussels (formerly *B. brevior*) showed that their free-living symbionts are present in biofilms, venting fluids as well as seawater away from hydrothermal activity (Harmer et al. 2008; Fontanez and Cavanaugh 2014). Our analyses for the mollusk symbioses confirm these findings but suggest that the distribution of free-living symbionts is not homogeneous across vent localities and habitat types. Whereas the Epsilon-like *Sulfurimonas* symbionts associated with *A. boucheti* were predominantly detected in biofilms at the central vent fields and showed increasing abundance in hydrothermal fluids towards the north, the Gamma1-like symbionts associated with *A. kojimai* and *A. strummeri* were most often found in biofilms and hydrothermal fluids at the southernmost vent field. By contrast, the *Bathymodiolus* symbionts were abundant in peripheral seawater and hydrothermal fluids at all vent fields, while the *Ifremeria* symbiont showed low but similar abundances in biofilms across the ELSC. These observations reflect both the regional distributions of *Alviniconcha*, *Ifremeria* and *Bathymodiolus* holobionts before the eruption as well as the local-scale habitat associations of these symbioses within single vent fields (Podowski et al. 2010; Beinart et al. 2012; Sen et al. 2013), highlighting how symbiont ecological niche influences host biogeography.

Our findings have important ramifications for the conservation biology of hydrothermal vent symbioses in the Lau Basin. The presence of gene flow among host populations as well as the detection of symbionts in the environment suggest that host species might be able to adapt and potentially recolonize disturbed vent habitats, although patterns of continuing decline in effective population size are alarming. In addition, there are multiple interacting factors that will determine repopulation potential and recruitment success at impacted vent fields. For example, northern ELSC vent fields were mostly devoid of hard substrate as they were buried under very thick ash deposits (Beinart et al. 2024). If the ash persists and is not removed via currents or other mechanisms, it is unclear whether soft substrate conditions will allow arriving larvae to settle successfully (Beinart et al. 2024). It is also unknown how temperature anomalies and changes in seafloor shape associated with the eruption (Clare et al. 2023; Seabrook et al. 2023) might alter ocean circulation patterns in the region, possibly impeding larval dispersal among vent fields. Increased fluid turbidity in these areas might further limit survival of host organisms by clogging gill filaments that are essential for respiration. This might be especially true for the *Alviniconcha* and *Ifremeria* gastropods which require access to higher hydrothermal fluid flows than the *Bathymodiolus* mussels (Podowski et al. 2010). Monitoring of recolonization during subsequent research cruises will provide valuable insights into the recovery process of hydrothermal vent populations after a natural disturbance and will advance our understanding of their resilience to anticipated human impacts from deep-sea mining (Civil Society Forum of Tonga 2024). Knowledge about these processes will be critical to develop sustainable management plans for the protection of these unique ecosystems.

## Acknowledgements

We thank the captains, crews and ROV pilots of the R/V *Thomas G. Thompson* (ROV *Jason II*) and R/V *Falkor* (ROV *Ropos*) for their support of the sample collections as well as the Kingdom of Tonga for allowing us access to their national waters and resources. We further thank Christine Li, Abi Goodman, Aidan Boving and Oliver Carey for helping with DNA extractions, Psomagen, Inc. for preparing and sequencing the metagenomic and RNA-seq libraries, and Argonne National Laboratory for preparing and sequencing the 16S amplicon libraries. This study was funded by the Schmidt Ocean Institute and the U.S. National Science Foundation (OCE-1736932 to R.A.B., OCE-0732369 to P.R.G. and EPSCoR Cooperative Agreement OIA-#1655221).

## Conflict of Interest

The authors declare no conflict of interest.

## Data Availability

Raw metagenomic data, 16S rRNA amplicon data and RNA-seq data have been deposited into NCBI’s Sequence Read Archive under BioProject numbers PRJNA1159215, PRJNA1157317, and PRJNA1157831. New *Alviniconcha and Ifremeria* transcriptomes have been submitted to the Transcriptome Shotgun Assembly database under accession numbers GKYL00000000, GKYM00000000, GKYP01000000 and GKYQ01000000, while symbiont MAGs have been submitted to GenBank under accession numbers listed in Supplementary Table S4. Bioinformatic code used to conduct analyses is available on GitHub (https://github.com/cbreusing/Lau_Basin_population_bottleneck).

## Notes

### Competing Interest Statement

The authors have declared no competing interest.

