## Supplementary Information for "Contrasting genomic responses of hydrothermal vent animals and their symbionts to population decline after the Hunga volcanic eruption"

**Supplementary Methods**

*Phenol:chloroform extraction protocol for SuPR and UFO filters*

This protocol outlines the steps required to extract DNA from water samples collected with the SuPR and UFO devices. Reagent amounts are provided for the extraction of five samples (four samples plus one extraction blank).

1. Prepare 400 mL DNA extraction buffer in a 1 L bottle by combining the following:

- 350 mL nuclease free water
- 4.85 g Tris Base
- 11.69 g EDTA
- 2.76 g NaH_2_PO_4_ * H_2_0
- 7.01 g NaCl

2. Stir the solution and adjust pH to 7.8–8 with NaOH. Bring the volume to 400 mL by adding nuclease free water. Autoclave the buffer, aliquot and freeze at -20°C.
3. Fill a 15 mL polypropylene tube for each sample with 1 mL silica/zirconia beads. Use 0.5 mm and 0.1 mm beads in a 4:1 ratio. Assemble scissors and tweezers for each sample and autoclave together with the tubes.
4. Sterilize a laminar flow hood by spraying the surface with ethanol. Cut open a large whirlpack and place it into the hood as sterile surface. Spray the whirlpack with ethanol. Place polypropylene tubes, scissors and tweezers into the hood and UV for at least 5 minutes.
5. Create lysozyme solution in a microcentrifuge tube. For five samples dissolve 0.035 g lysozyme in 8.4 µL Tris-HCl and top it off to 700 µL with nuclease-free water. Dissolve at the beginning of the extraction day and leave in the fridge.
6. Take samples out of the -80°C freezer and let thaw briefly. Cut each filter into small pieces (<1 cm^2^) and place in an empty, labeled 50 mL tube.
7. Add 7 mL of extraction buffer to the filter using a sterile plastic serological pipette.
8. Add 140 µL of lysozyme to the filter using a regular pipette.

**The following steps need to be performed under a fume hood.**

9. Under the fume hood make 1 M dithiothreitol (DTT). For five samples add 0.2692 g DTT to 1750 µL nuclease free water. Flick the DTT. Add 350 µL of 1M DTT to each sample tube.
10. Wrap the 50 mL sample tubes with parafilm to prevent leakage. Vortex for ~1 minute and carefully invert the tube multiple times to soak the sample filter with buffer.
11. Incubate on a rotator at 37°C for 30 minutes. Check sample from time to time for leakage.
12. After 30 minutes remove samples and adjust temperature to 65°C.
13. Under the fume hood tap tubes on counter to remove filter pieces from caps. Add 180 µL proteinase K and 350 µL 20% SDS to each sample.
14. Use a sterile pipette tip to remove filter pieces from tube caps. Wrap tightly with parafilm. Vortex and invert to mix.
15. Incubate samples on a rotator at 65°C for 45 minutes (or up to 1.5 hours). Check frequently for leakage!
16. After incubation, pour as much of the ~8 mL (ideally 6 mL) supernatant from the 50 mL sample tubes into the 15 mL bead tubes. Be careful not to transfer any filter material. Do not fill past the 7 mL line.
17. Ensure cap is tight on the 15 mL tubes and no filter or beads are blocking the seal. Vortex on high for 10 minutes or homogenize in a bead beater.
18. Incubate on ice for 5-10 minutes. Centrifuge at 2500 g for 5-10 minutes.
19. While waiting, fill a 50 mL tube with 100% ethanol and incubate at -20°C.
20. Take samples out of the centrifuge and decant supernatant into new 15 mL tube. Be careful not to dislodge the pellet.
21. Put on a second pair of gloves and protective goggles.
22. Under the fume hood, add an equal volume of 25:24:1 phenol:chloroform:isoamyl alcohol to the sample using a 10 mL glass pipette. Close the tube and make sure the cap is on tight. Wipe the tube with a Kimwipe. Shake for 1 minute or vortex for 10 seconds.
23. Centrifuge at 2500 g for 10 minutes. If phase separation is weak, repeat this step.
24. Remove as much of the top aqueous phase as possible and transfer into new 15 mL tube. Be careful not to touch the white middle layer containing organic inhibitors.
25. Add an equal volume of 24:1 chloroform:isoamyl alcohol to the aqueous phase using a 10 mL glass pipette.
26. Shake for 1 minute or vortex for 10 seconds.
27. Centrifuge at 2500 g for 10 minutes. Take 20 mg/mL glycogen out of the -20°C freezer to thaw.
28. Remove as much of the top aqueous phase as possible and transfer into a new, labeled 50 mL Falcon tube. Be careful not to touch the white middle layer.
29. Add 2 volumes of 100% ethanol from the -20°C freezer and 1:50 volume of 20 mg/mL glycogen. Make sure tube is sealed tight and then mix carefully without letting liquid touch the cap.
30. Incubate samples at -20°C for 0.5-2 hours or overnight. Alternatively, incubate samples at -80°C for <30 minutes without letting samples freeze solid. Check samples every 10 minutes if incubating at -80°C.
31. Centrifuge at 15,000 g for 30 minutes to pellet DNA.
32. Prepare 50 mL of 70% ethanol and store at -20°C.
33. Remove supernatant from the sample tubes without disturbing the pellet.
34. Gently add 5-10 mL (~8 mL) of cold 70% ethanol to wash the pellet. Gently roll the tube. Do not disturb the pellet.
35. Centrifuge at 15000 g for 30 minutes. Set water bath or incubator to 50°C.
36. Remove supernatant and allow pelleted DNA to air dry for 30-40 minutes or until no traces of ethanol can be seen. If pellets are stable, let air dry by placing tubes upside down on a tube rack.
37. Resuspend pellet in 100 µL of nuclease free water and incubate at 50°C for 2-10 minutes. Pipette up and down several times and/or scrape tube with pipette tip.
38. Purify samples with the Qiagen DNeasy PowerClean Pro Cleanup Kit.
39. Quantify final extracts with the Qubit dsDNA HS Assay Kit and store at -80°C.

**Supplementary Figures**

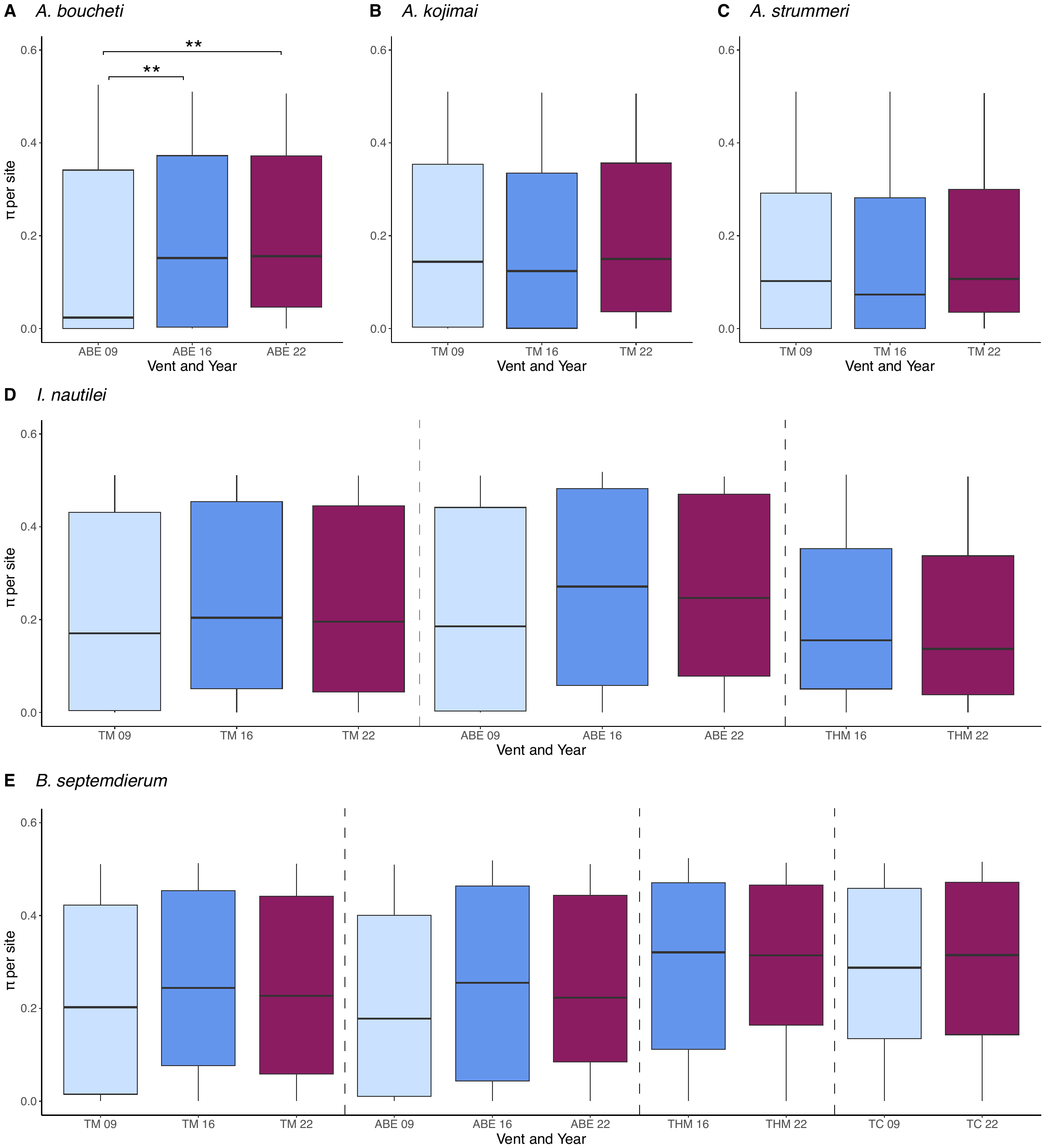

**Fig. S1** Bar plots of per-site nucleotide diversities at high-quality variants in host populations across years at selected vent sites. Stars indicate significant differences among groups. * *p* < 0.05, ** *p* < 0.01, *** *p* < 0.001

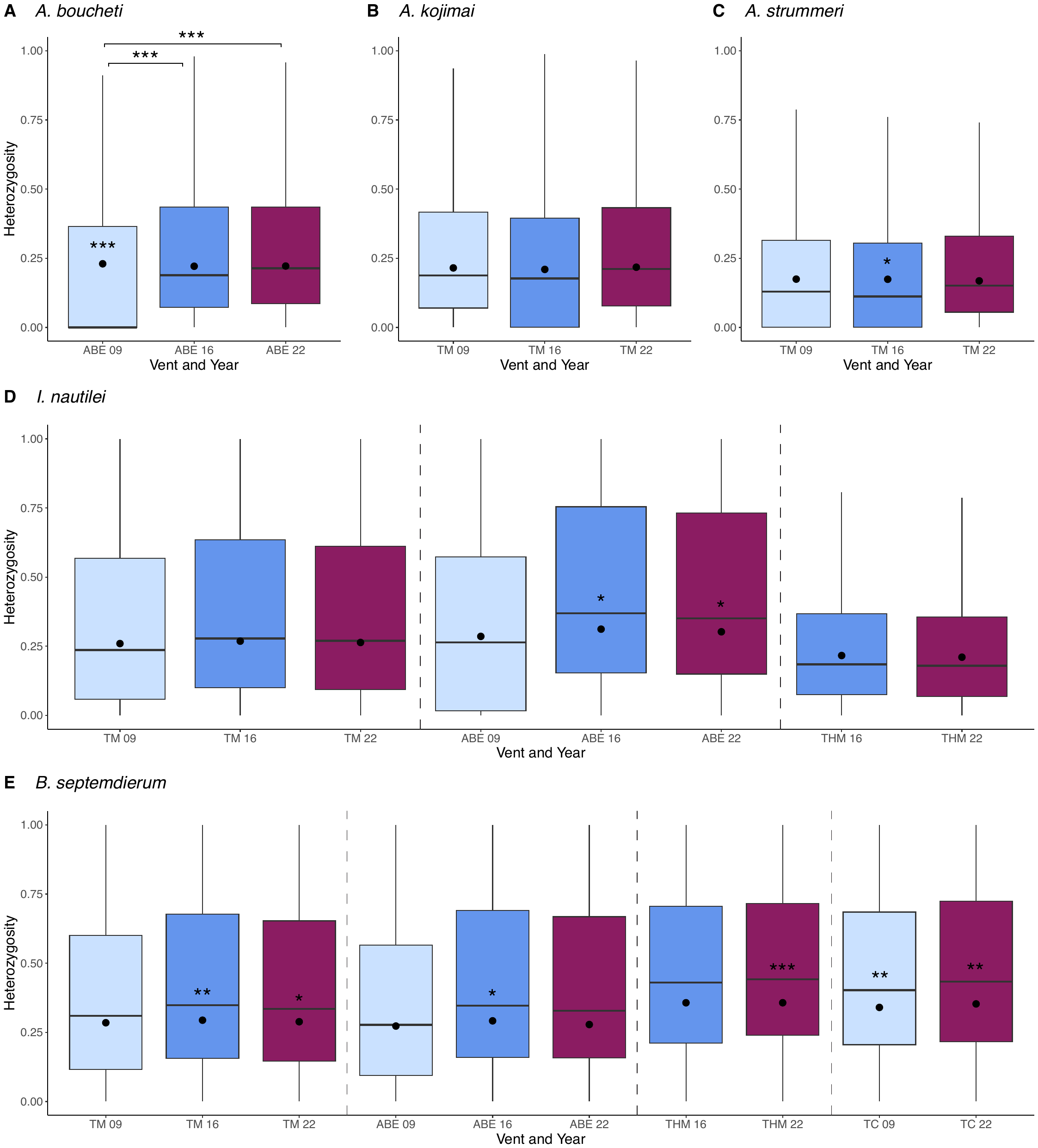

**Fig. S2** Bar plots of observed heterozygosities at high-quality variants in host populations across years at selected vent sites. Black dots indicate median expected heterozygosities. Stars within bar plots show significant differences between expected and observed mean heterozygosities for a group, while stars above bar plots denote significant differences in mean observed heterozygosities between groups. * *p* < 0.05, ** *p* < 0.01, *** *p* < 0.001

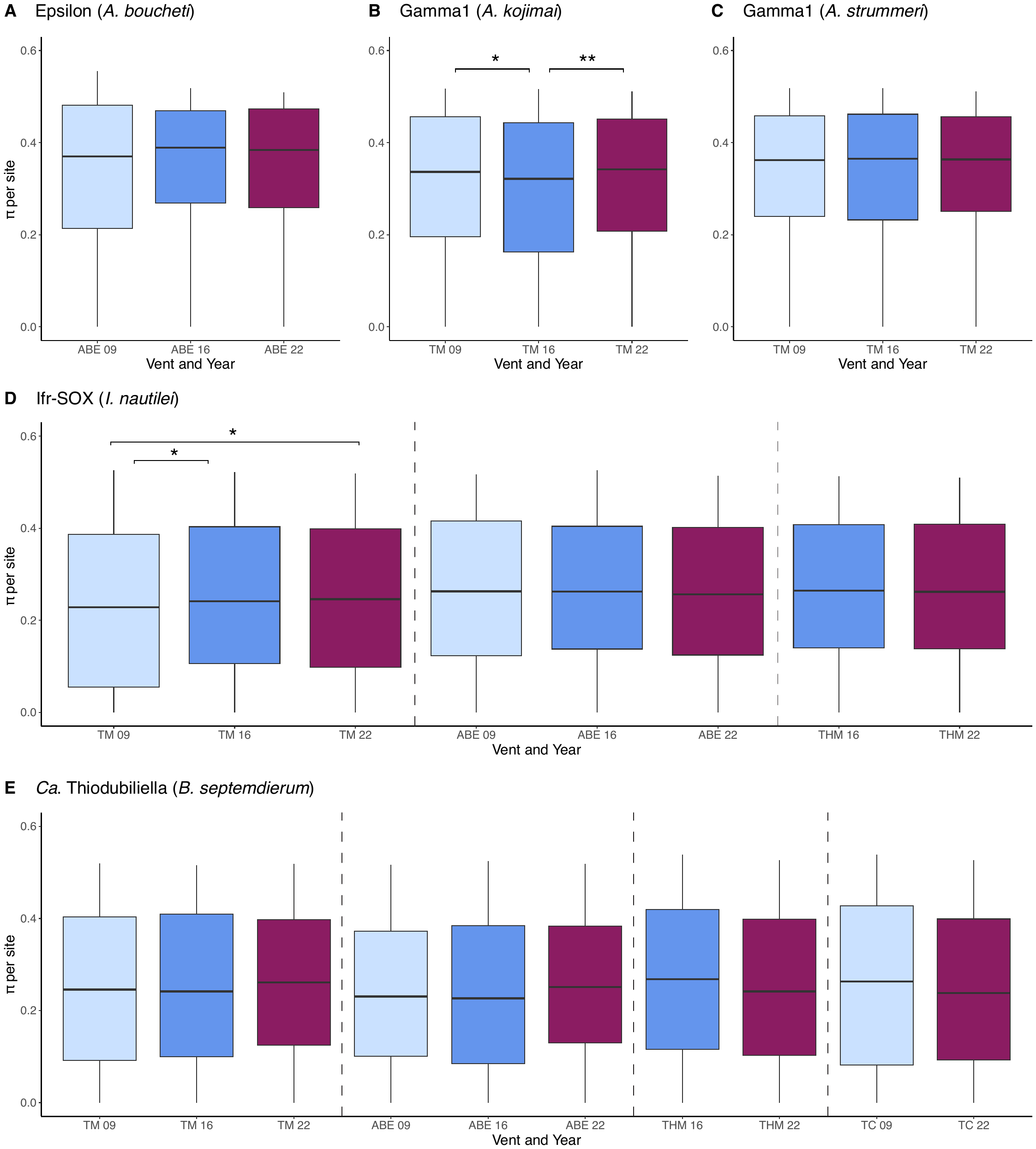

**Fig. S3** Bar plots of per-site nucleotide diversities at high-quality variants in symbiont populations across years at selected vent sites. Stars indicate significant differences among groups. * *p* < 0.05, ** *p* < 0.01, *** *p* < 0.001

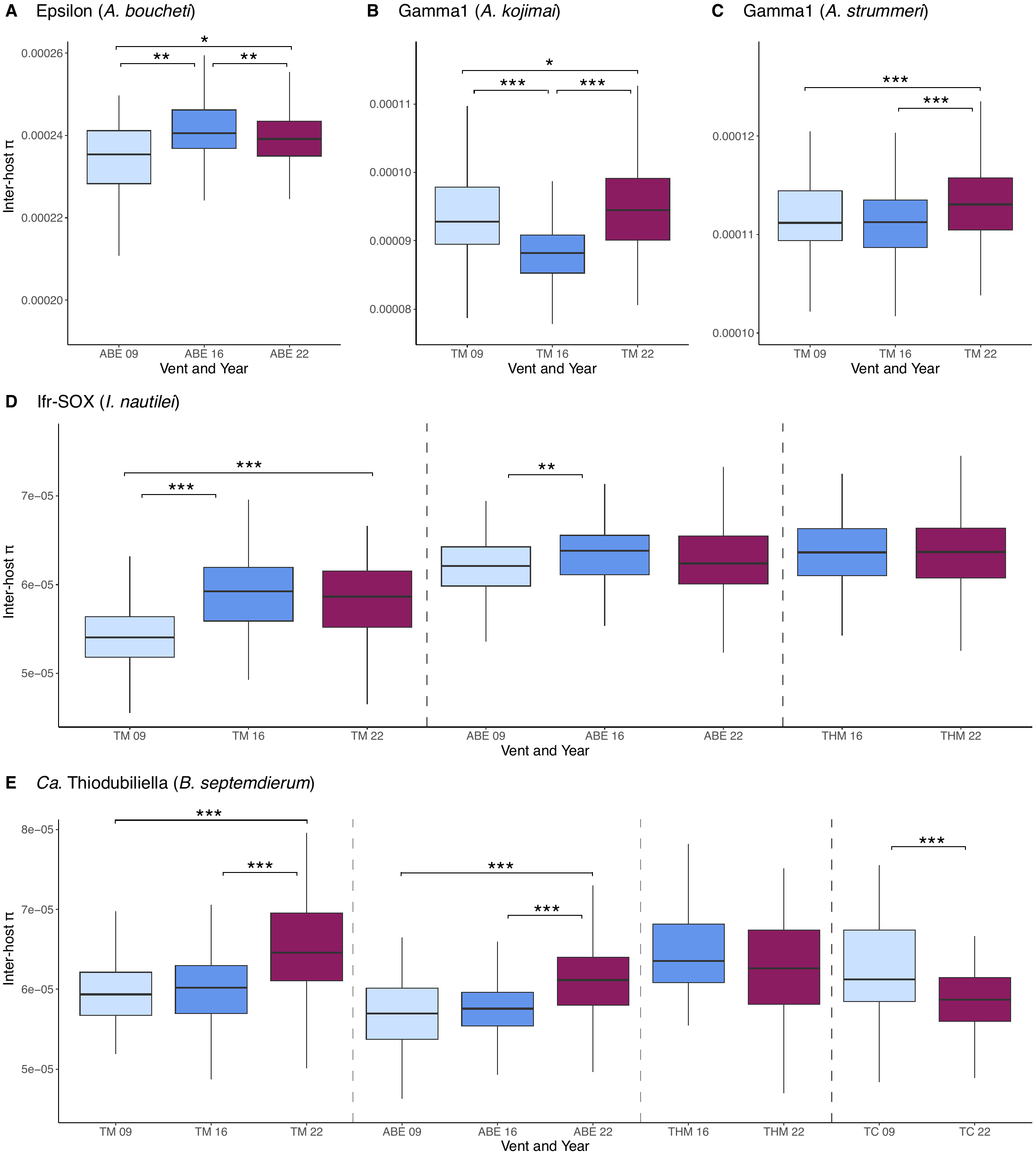

**Fig. S4** Bar plots of inter-host diversities between symbiont populations across years at selected vent sites. Stars indicate significant differences among groups. * *p* < 0.05, ** *p* < 0.01, *** *p* < 0.001

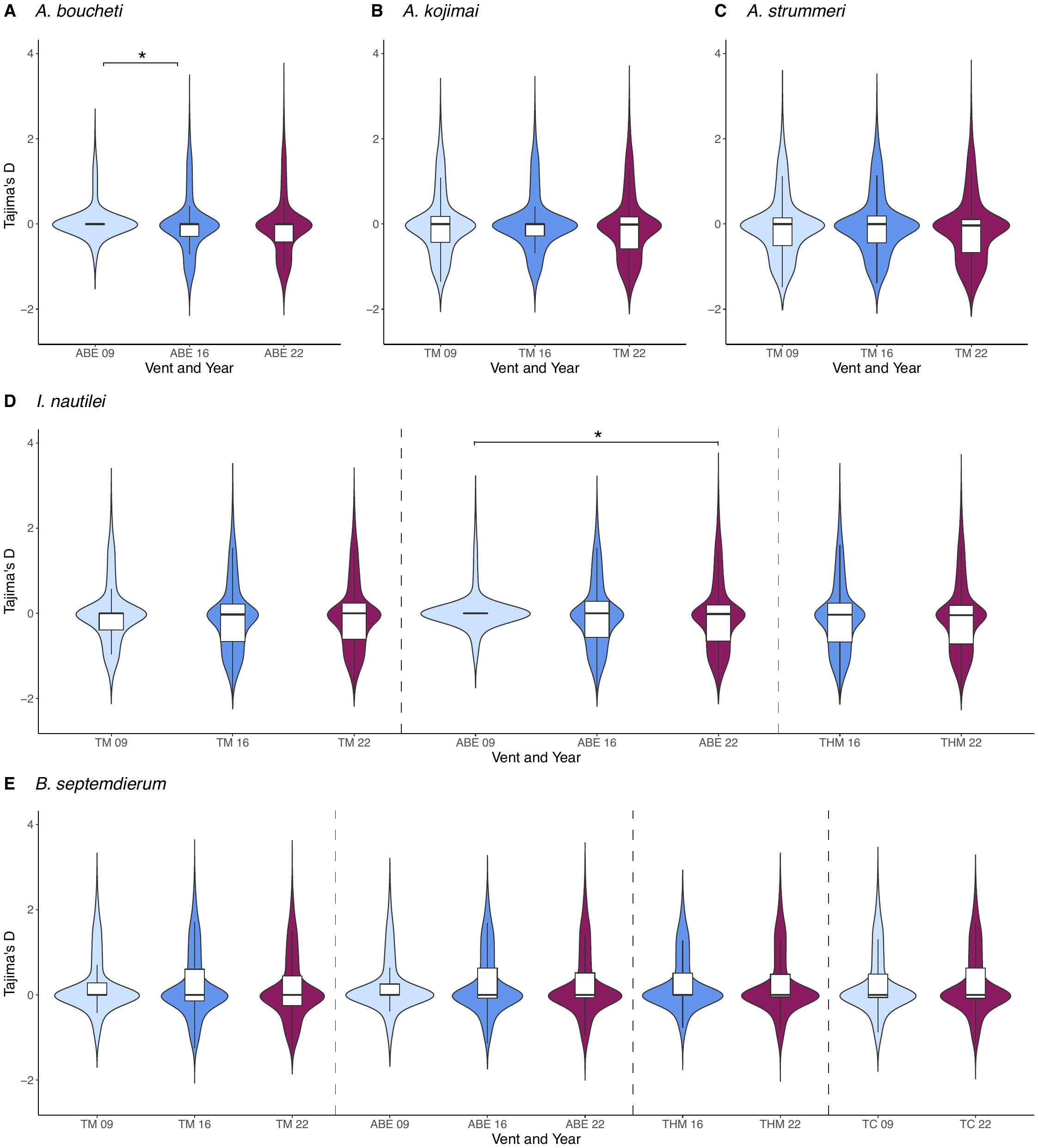

**Fig. S5** Violin plots with integrated bar plots for Tajima’s D values calculated across host transcriptomes between years at selected vent sites. Stars indicate significant differences among groups. * *p* < 0.05, ** *p* < 0.01, *** *p* < 0.001

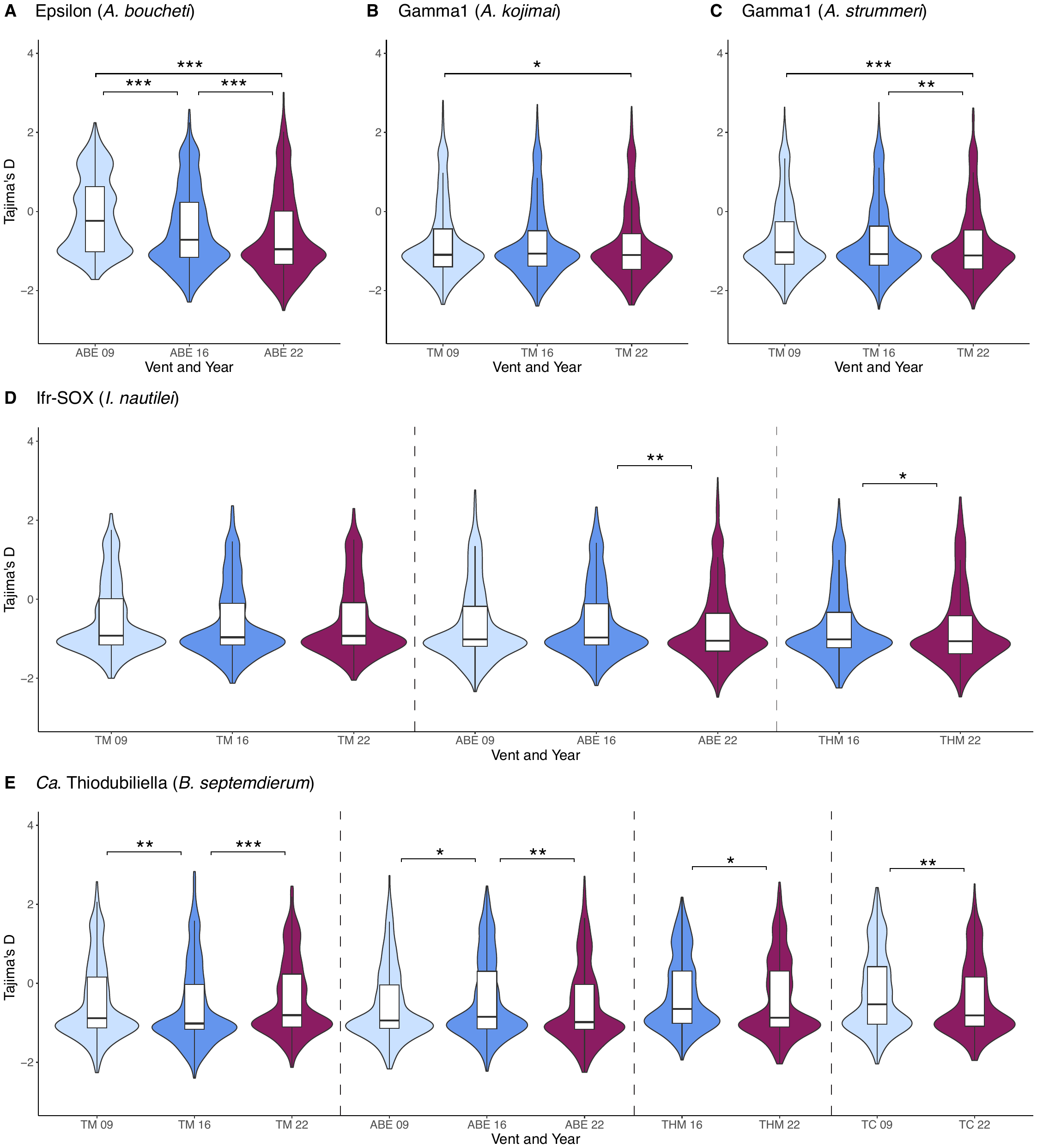

**Fig. S6** Violin plots with integrated bar plots for Tajima’s D values calculated across symbiont pangenomes between years at selected vent sites. Stars indicate significant differences among groups. * *p* < 0.05, ** *p* < 0.01, *** *p* < 0.001
