## Supplementary figures and images for "Contrasting genomic responses of hydrothermal vent animals and their symbionts to population decline after the Hunga volcanic eruption"

### Supplementary Figure S1

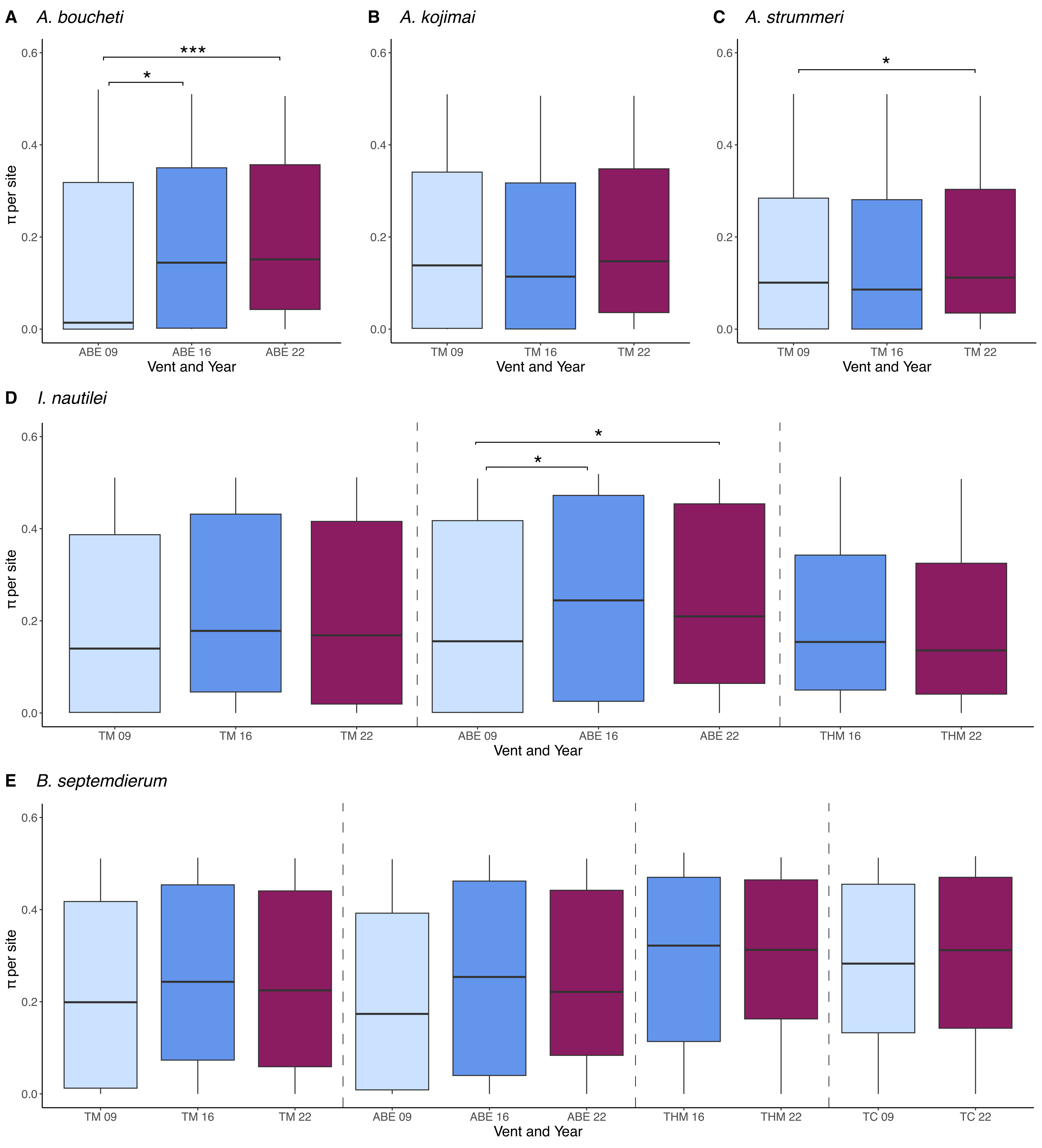

### Supplementary Figure S2

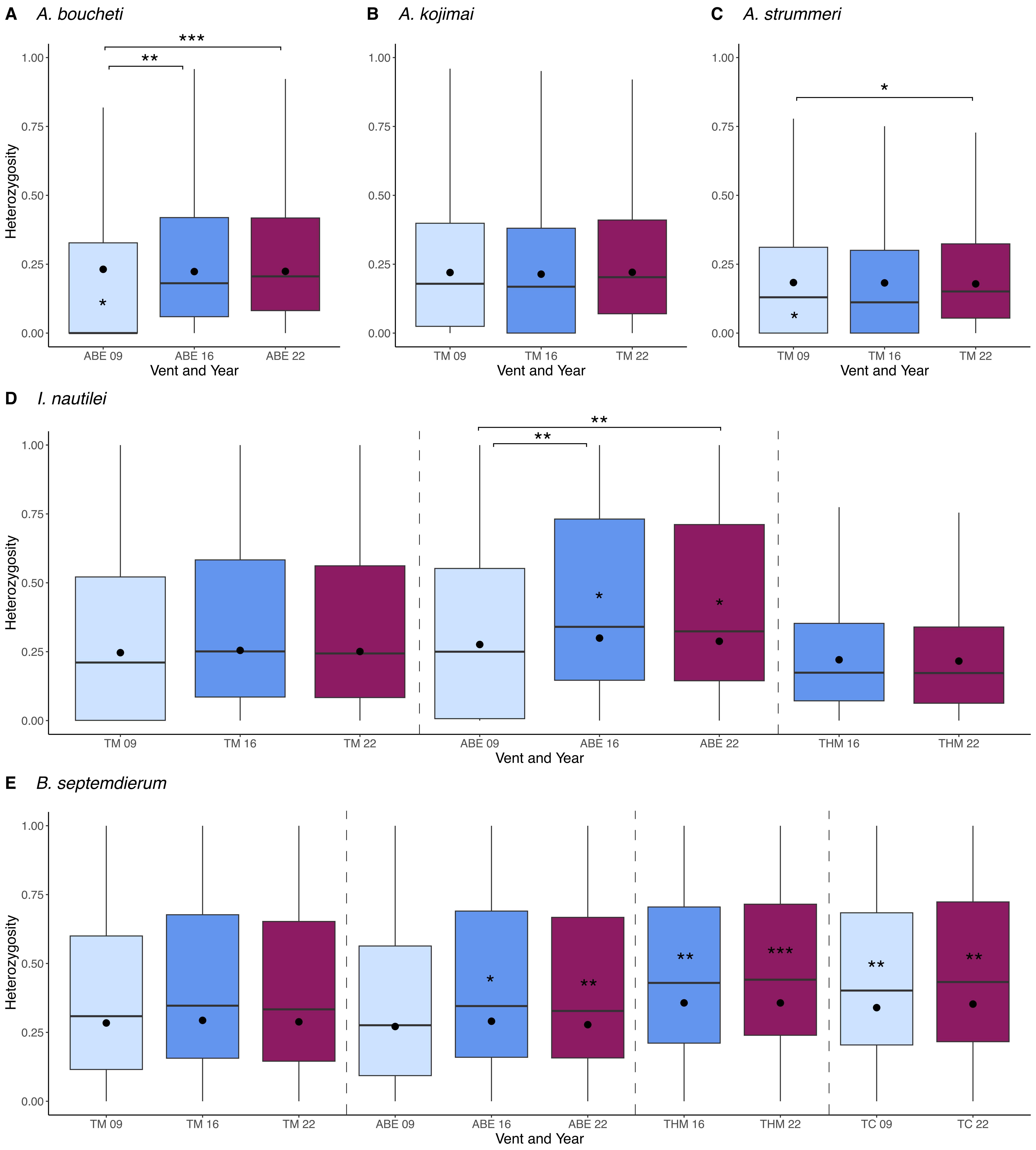

### Supplementary Figure S3

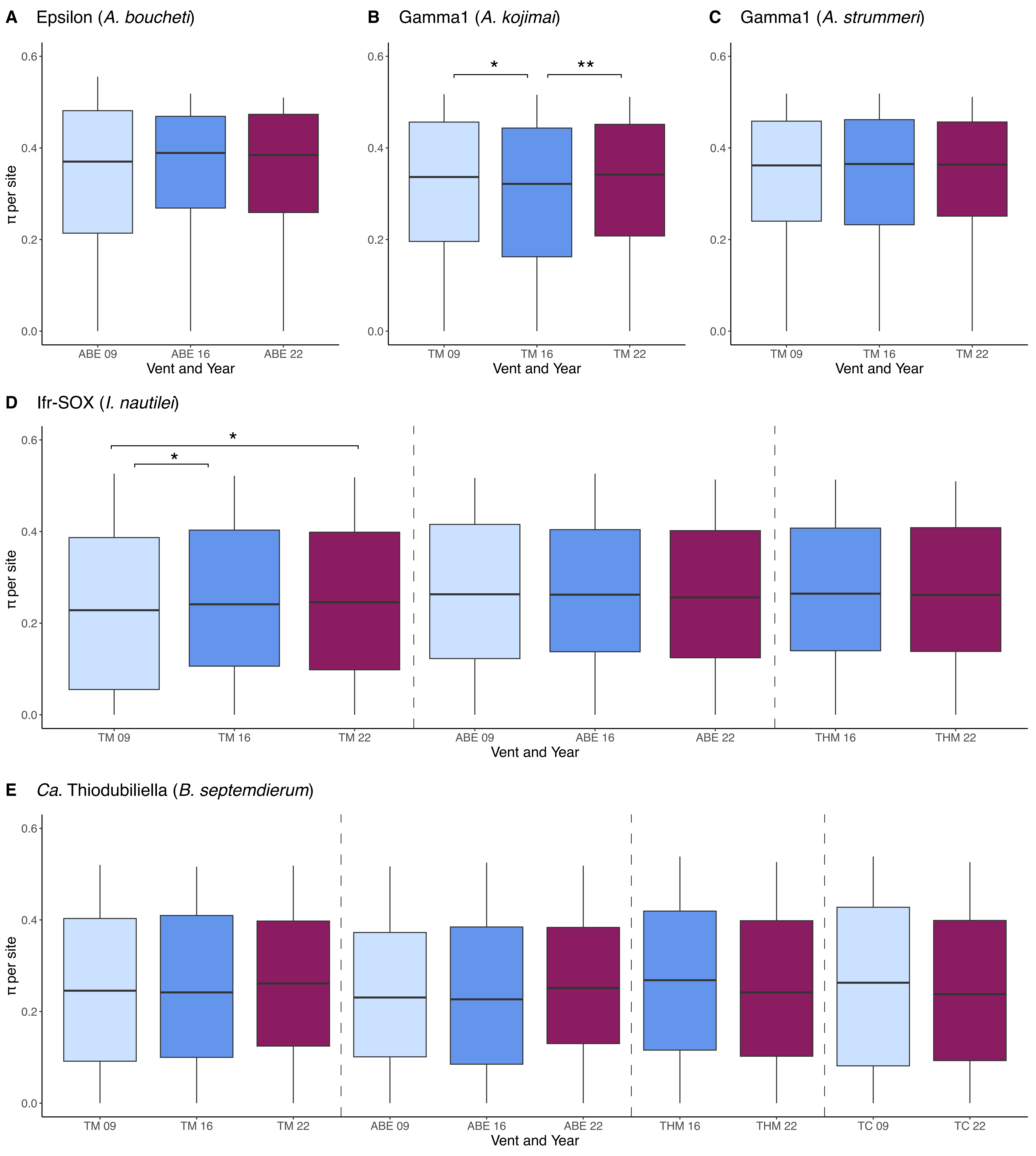

### Supplementary Figure S4

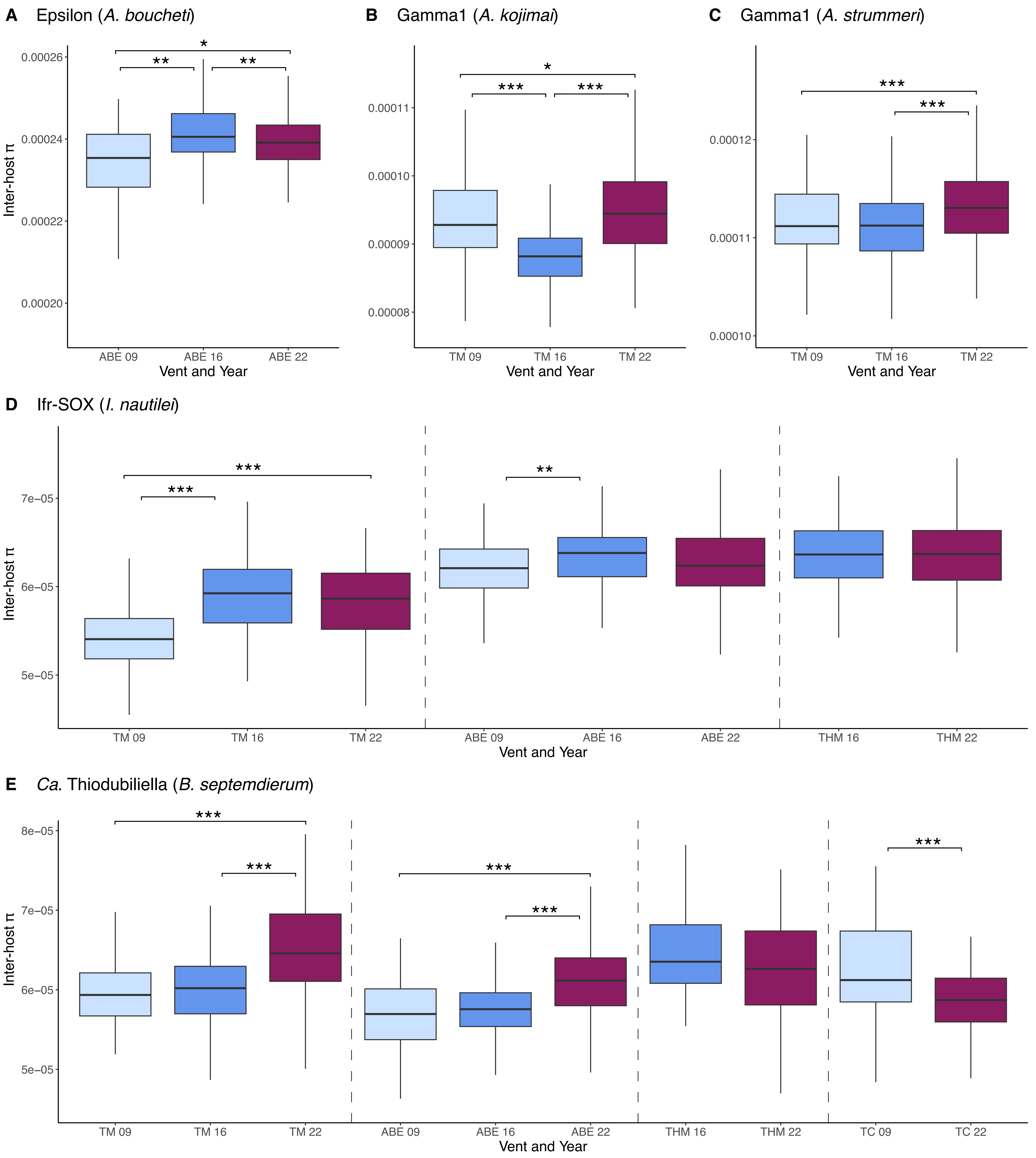

### Supplementary Figure S5

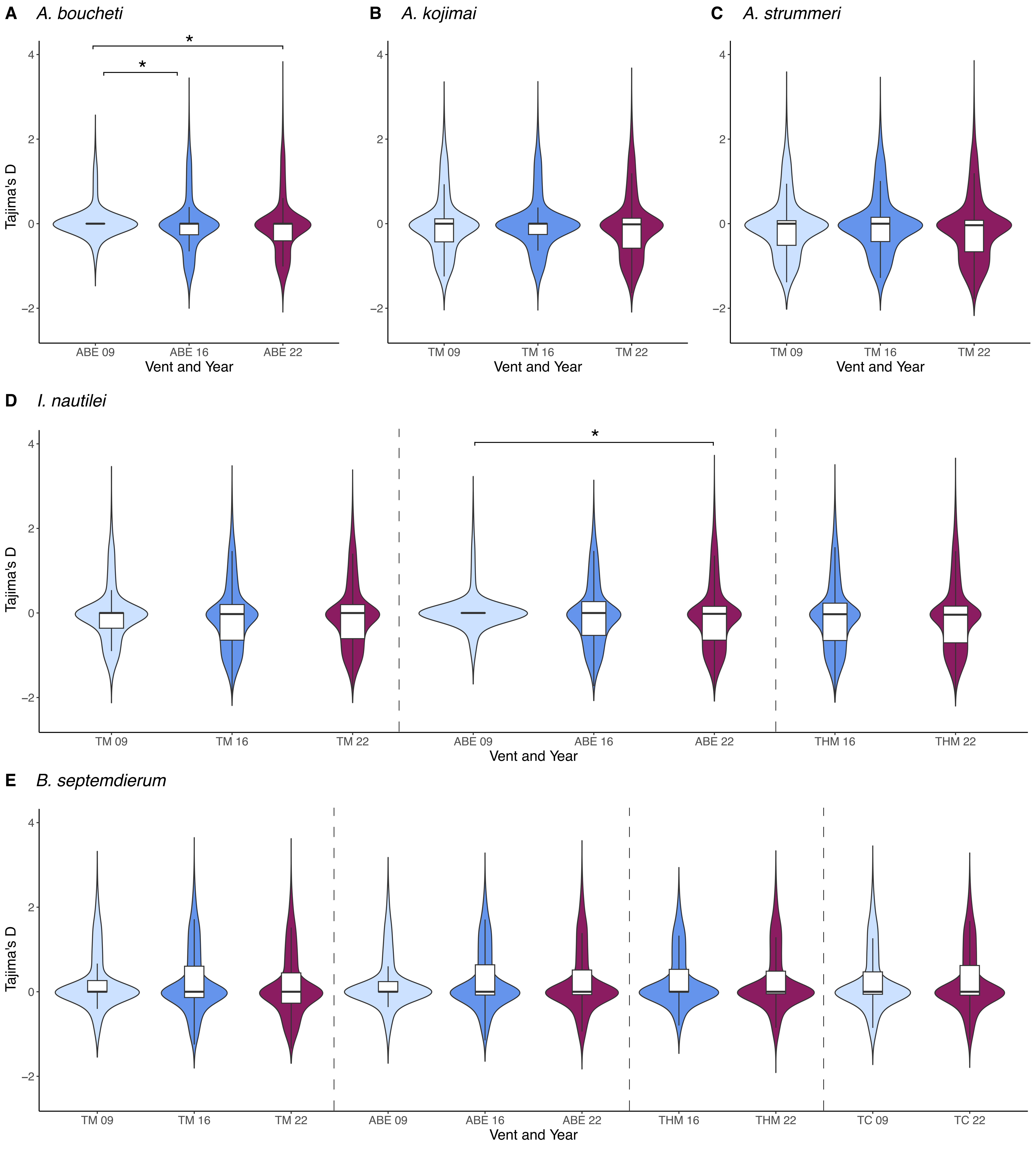

### Supplementary Figure S6

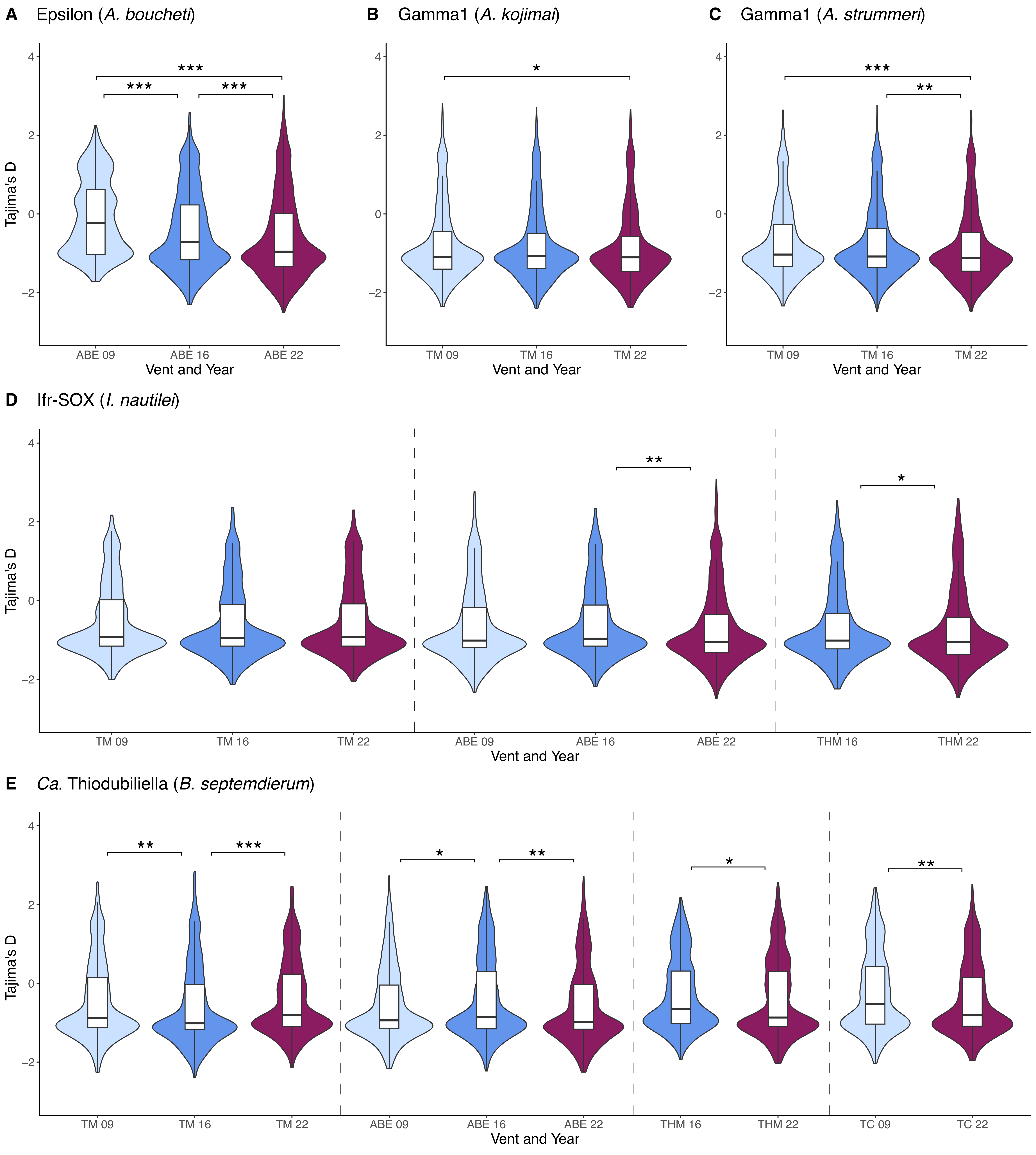
